# Functional significance of PUF partnerships in *C. elegans* germline stem cells

**DOI:** 10.1101/2023.02.15.528708

**Authors:** Ahlan S. Ferdous, Stephany J. Costa Dos Santos, Charlotte R. Kanzler, Heaji Shin, Brian H. Carrick, Sarah L. Crittenden, Marvin Wickens, Judith Kimble

## Abstract

PUF RNA-binding proteins are conserved stem cell regulators. Four PUF proteins govern self-renewal of *C. elegans* germline stem cells together with two intrinsically disordered proteins, LST-1 and SYGL-1. Based on yeast two-hybrid results, we proposed a composite self-renewal hub in the stem cell regulatory network, with eight PUF partnerships and extensive redundancy. Here, we investigate LST-1–PUF and SYGL-1–PUF partnerships and their molecular activities in their natural context – nematode stem cells. We confirm LST-1–PUF partnerships and their specificity to self-renewal PUFs by co-immunoprecipitation and show that an LST-1(A^m^B^m^) mutant defective for PUF-interacting motifs does not complex with PUFs in nematodes. LST-1(A^m^B^m^) is used to explore the functional significance of the LST-1–PUF partnership. Tethered LST-1 requires the partnership to repress expression of a reporter RNA, and LST-1 requires the partnership to co-immunoprecipitate with NTL-1/Not1 of the CCR4-NOT complex. We suggest that the partnership provides multiple molecular interactions that work together to form an effector complex on PUF target RNAs. Comparison of PUF-LST-1 and Pumilio–Nanos reveals fundamental molecular differences, making PUF–LST-1 a distinct paradigm for PUF partnerships.

**Summary statement:** Partnerships between PUF RNA-binding proteins and intrinsically disordered proteins are essential for stem cell maintenance and RNA repression.

## Introduction

RNA-binding proteins are central to gene regulation and a wide range of biological phenomena and human diseases (Gebauer et al., 2021; Gong et al., 2022; Matia-González et al., 2015). Often, they work within regulatory complexes that modulate their activity. Most relevant to this work, PUF RNA-binding proteins (PUF for <u>Pu</u>milio and <u>F</u>BF) are broadly conserved regulators of gene expression. From yeast to humans, PUF proteins bind mRNAs with exquisite sequence specificity, and repress RNA stability or translation (Goldstrohm et al., 2018; Miller & Olivas, 2011; Wickens et al., 2002; Zamore et al., 1997). Moreover, PUF proteins have conserved biological roles in stem cells and neurobiology, with recently discovered links to human disease (Gennarino et al., 2018; Gong et al., 2022; Naudin et al., 2017; Rajasekaran et al., 2022). Great progress has been made understanding the molecular function of PUF proteins themselves, but PUFs interact with numerous other proteins (Campbell et al., 2012; Friend et al., 2012; Ginter-Matuszewska et al., 2011; Jaruzelska et al., 2003; Moore et al., 2003; Wu et al., 2013). Partnership with Nanos, for example, enhances PUF binding affinity to RNA and refines its recognition sequence (Weidmann et al., 2016). The functions of other partnerships, however, are poorly characterized and represent the next frontier in understanding how PUF proteins control gene expression.

PUF-partner complexes have been implicated in self-renewal of germline stem cells (GSC) in the nematode *C. elegans* (Haupt et al., 2020; Shin et al., 2017). These PUF partnerships form a composite node in the GSC regulatory network, dubbed the “PUF hub” (Figure 1A). Molecular evidence for eight PUF partnerships was based on assays in yeast *or in vitro,* all done with incomplete protein fragments (Haupt et al., 2020; Qiu et al., 2019; Qiu et al., 2022; Shin et al., 2017). While those experiments were powerful, a deep understanding of how PUF proteins regulate RNAs in stem cells demands testing their molecular activities in the cells where they normally act. In this work, we investigate PUF hub partnerships *in their natural context – C elegans* germline stem cells – and do so for full-length proteins with validated biological functions. As an introduction to this complex mesh of regulators, we first describe the key PUF proteins, then the partners and finally their partnerships.

Four PUF proteins belong to the self-renewal hub (Crittenden et al., 2002; Haupt et al., 2020) (Figure 1A). FBF-1 and FBF-2 are nearly identical to each other and play the more prominent role; PUF-3 and PUF-11 also have sequences similar to each other but play a more minor role. Like other PUF proteins, the four self-renewal PUFs bind to sequence elements in the 3’UTR of their target mRNAs (Hubstenberger et al., 2012; Koh et al., 2009; Zhang et al., 1997) and are best known for repression of differentiation RNAs in the GSC pool (Crittenden et al., 2002; Merritt et al., 2008). The four PUFs are variably redundant with each other: no major GSC defect occurs in any of the single mutants (*fbf-1, fbf-2, puf-3* or *puf-11*) or in the *puf-3 puf-11* double mutant. However, all GSCs are lost to differentiation at the last larval stage in *fbf-1 fbf-2* double mutants, and in early larvae of *fbf-1 fbf-2; puf-3 puf-11* quadruple mutants. Thus, these four PUF proteins are responsible for GSC self-renewal throughout development.

Two PUF partners, LST-1 and SYGL-1, also belong to the hub (Haupt et al., 2020; Kershner et al., 2014; Shin et al., 2017) (Figure 1A). Both proteins are composed largely of regions of low complexity, which are predicted to be intrinsically disordered (IDRs). The two proteins bear no sequence similarity but they are functionally redundant: no major GSC defect occurs in either single mutant *(lst-1* or *sygl-1)*, but all GSCs are lost in early larvae of *lst-1 sygl-1* double mutants (Kershner et al., 2014). Consistent with a key role in self-renewal, both proteins are restricted to the GSC pool, and expanded expression of either LST-1 or SYGL-1 drives formation of a germline tumor (Shin et al., 2017). Thus, the stem cell function of LST-1 and SYGL-1 is equivalent to that of the self-renewal PUFs – they are responsible for GSC self-renewal throughout development.

**Figure 1:**
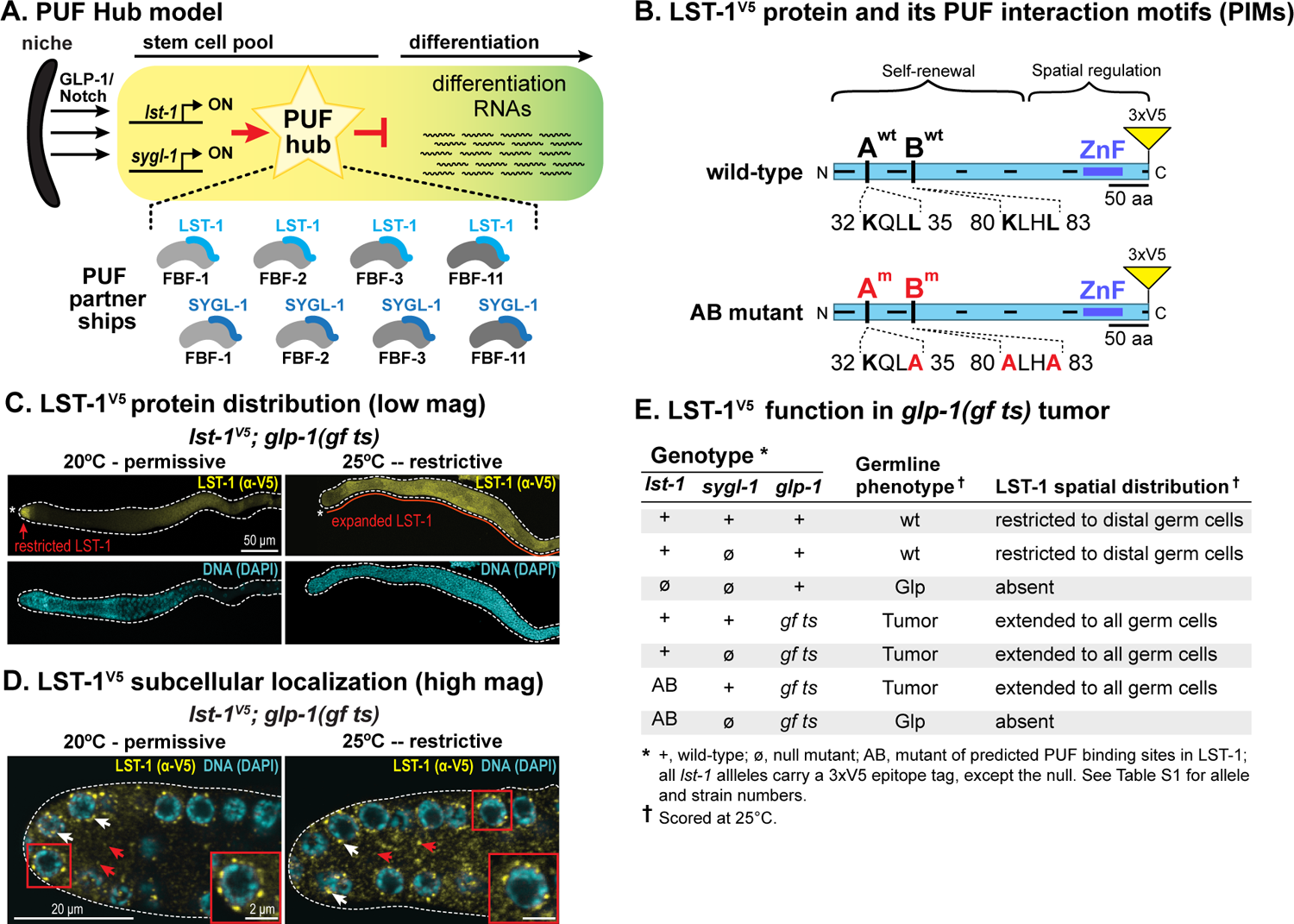
PUF partnerships in the PUF hub and their biochemical analysis in nematodes. A. PUF hub model. LST-1 and SYGL-1 are central to a composite regulatory hub: each is proposed to partner with any of four PUF proteins (FBF-1, FBF-2, PUF-3 and PUF-11, grey) and to repress differentiation RNAs for maintenance of the GSC pool. GLP-1/Notch signaling activates *lst-1* and *sygl-1* transcription (black arrows) at the distal end of the gonad, which restricts LST-1 and SYGL-1 expression to the GSCs. B. LST-1 protein architecture. LST-1 possesses an N-terminal “self-renewal” domain composed of multiple intrinsically disordered regions (IDRs, black lines along protein axis) and a C-terminal “spatial regulation” domain with additional IDRs and a Zinc finger (ultramarine blue). Within the self-renewal region are two PUF interaction motifs (A and B), shown with wild-type (top) and mutant (bottom) sequences. C. LST-1^V5^ distribution expands in *glp-l(gf ts)* mutants. Representative confocal z-projections of extruded gonads stained with α-V5 antibody to detect LST-1^V5^ (yellow) and with DAPI (cyan) for DNA. Left panel, red arrow marks spatially restricted LST-1^V5^ at permissive temperature, as in wild-type (Haupt et al., 2019; Shin et al., 2017); right panel, red line marks expanded LST-1^V5^ in a germline tumor, formed at restrictive temperature. Asterisk indicates distal end of the gonad, and dotted line marks its boundary. D. Subcellular distribution of LST-1^V5^ in *glp-l(gf ts)* germline. Representative images of single confocal z-slices from middle plane of distal region of extruded gonads stained with α-V5 antibody to detect tagged LST-1^V5^ (yellow). LST-1^V5^ exists in both perinuclear puncta (white arrows) and the cytoplasm (red arrows). Inset shows higher magnification. E. LST-1^V5^ retains stem cell regulatory function when expanded in *glp-1(gf ts)* germline tumors.

The first clue that LST-1 and SYGL-1 might function together with PUF proteins in a complex came from a genetic finding that forced overexpression of LST-1 or SYGL-1 could not form a tumor in the absence of FBF-1 and FBF-2 (Shin et al., 2017). Why might LST-1 and SYGL-1 depend on these two PUF proteins? LST-1 and SYGL-1 interacted with FBF-1 and FBF-2 in yeast two-hybrid assays (Shin et al., 2017), and similarly interacted with PUF-3 and PUF-11 (Boxem et al., 2008; Haupt et al., 2020; Qiu et al., 2019; Racher & Hansen, 2012). These findings crystallized the idea that LST-1 and SYGL-1 likely function as PUF partners. Consistent with that idea, an FBF target RNA, *gld-1,* was de-repressed in *lst-1 sygl-1* double mutants (Shin et al., 2017).

More recently, two PUF-interacting motifs (PIMs) were identified in the LST-1 amino acid sequence (Figure 1B) (Haupt et al., 2019). The “KxxL” sequence of the LST-1 PIMs was similar to the “KTxL” PIMs in other FBF partners, GLD-3 and CPB-1 (Campbell et al., 2012; Menichelli et al., 2013; Wu et al., 2013). By yeast two-hybrid, an LST-1 protein with only one intact PIM could still bind PUF, but binding was lost when both were mutated. The biological impact of each LST-1 PIM (PIM-A and PIM-B) was assayed in nematodes that lacked SYGL-1, its redundant counterpart. Here again, LST-1 protein mutated for a single PIM, LST-1(A^m^) or LST-1(B^m^), retained its ability to maintain germline stem cells, but a double PIM mutant, LST-1(A^m^B^m^), could not (Haupt et al., 2019). Indeed, the two PIMs reside within a 210 amino acid “self-renewal region” that harbors multiple IDRs and is both necessary and sufficient to maintain stem cells (Fig 1B)(Haupt et al 2019). *In vitro*, short LST-1 peptides carrying either of the two LST-1 PIMs bind to FBF-2 at the same site as CPB-1 and GLD-3 (Menichelli et al., 2013; Qiu et al., 2019; Qiu et al., 2022; Wu et al., 2013). Together, these findings suggested that LST-1–PUF partnerships are important for stem cell regulation.

These earlier studies set the stage for testing the PUF hub model in nematodes and analyzing the molecular activities of self-renewal PUF partnerships. Here, we confirm that LST-1 physically associates with self-renewal PUF proteins in nematodes, but that a mutant lacking the LST-1 PIMs, LST-1(A^m^B^m^), does not. The LST-1(A^m^B^m^) mutant thus provides an incisive and unique tool to probe the functional significance of PUF partnerships *in vivo*. We demonstrate that LST-1 possesses repressive activity when tethered to a reporter RNA, and that its PUF partnership is essential for repression. We show further that LST-1 must partner with a PUF to physically associate with the CCR4-Not complex (CNOT). Based on these findings, we propose that the LST-1–PUF partnership is responsible for multiple molecular interactions that together form a stable effector complex on PUF target RNAs. Finally, we provide evidence that SYGL-1 functions much like the LST-1 in GSC maintenance and RNA repression.

## Results

### LST-1 associates *in vivo* with PUF proteins integral to the self-renewal hub

LST-1 and SYGL-1 are expressed at low levels in whole worm extracts because of their spatial restriction to GSCs. We previously used a strong germline promoter to increase LST-1 and SYGL-1 abundance and managed to coIP SYGL-1 with a single PUF protein, FBF-2 (Shin et al., 2017). However, that approach was technically challenging; it could not be extended to other PUF proteins for SYGL-1 and was unsuccessful for LST-1. To probe LST-1–PUF partnerships *in vivo,* we sought a different way to increase LST-1 levels. A conditional mutant of the GLP-1/Notch receptor, *glp-1(ar202)*, causes constitutive Notch signaling at restrictive temperature (25°), expands the number of GSCs and drives formation of a germline tumor (Pepper et al., 2003); henceforth, we call this mutant *glp-1(gf ts)*. Because the *lst-1* gene is a direct target of GLP-1/Notch signaling (Kershner et al., 2014; Lee et al., 2016), we expected that constitutive–Notch signaling would increase LST-1 levels. We therefore generated a strain carrying *glp-l(gf ts)* and 3×V5 epitope-tagged LST-1. LST-1^V5^ was previously shown to retain wild-type LST-1 activity in stem cell regulation (Haupt et al., 2019). At the permissive temperature of 20°C, GSCs were maintained normally and LST-1^V5^ distribution appeared normal, but at restrictive temperature, a germline tumor formed and LST-1^V5^ expanded to fill that tumor (Figure 1C). As reported previously (Pepper et al., 2003), small patches of differentiating cells were sometimes seen in the tumors, and LST-1^V5^ was missing from those patches (Fig S1). Within germ cells, LST-1^V5^ was located as normal in perinuclear granules and cytoplasm at both temperatures (Figure 1D). We conclude that LST-1^V5^ retains normal activity but that its expression becomes abundant with a simple shift to restrictive temperature in this easily maintained strain.

To ask whether the LST-1–PUF interactions found in yeast reflect interactions in nematodes, we generated a set of strains for co-immunoprecipitations (coIPs). Each strain carried *glp-1(gf ts)* and distinctly tagged LST-1 and PUF proteins (*see Supplement for specific genotypes)*. Control strains included *glp-1 (gf ts)* and each PUF tagged allele. LST-1^V5^ and LST-1^FLAG^ both function normally in genetic assays (Figure 1E; Haupt et al., 2019), and similarly all tagged PUFs behave normally (Figure S2A). Moreover, all tagged PUFs were expressed throughout the germline at high levels in the tumors at restrictive temperature (Fig S2B).

For the co-immunoprecipitations, we prepared lysate from at least 10^6^ synchronized adults with germline tumors; all animals were cross-linked with formaldehyde prior to collection. At least two replicates of each IP had similar results, both here and for other IPs reported in this work. Among the PUF proteins in the self-renewal hub, LST-1 immunoprecipitation brought down FBF-1, FBF-2 and PUF-11 (Figure 2A-C); PUF-3 was least abundant and not attempted. In contrast to the self-renewal PUFs in the hub, LST-1 did not coIP with PUF-8, which is present in germline stem cells but is not essential for stem cell maintenance (Figure 2D). Consistent with that finding, key residues in FBF-2, critical for LST-1 binding, are conserved in all four self-renewal PUFs, but not in PUF-8 (Qiu et al., 2019; Wu et al., 2013). We conclude that LST-1 associates in the nematode specifically with PUF proteins in the hub.

**Figure 2:**
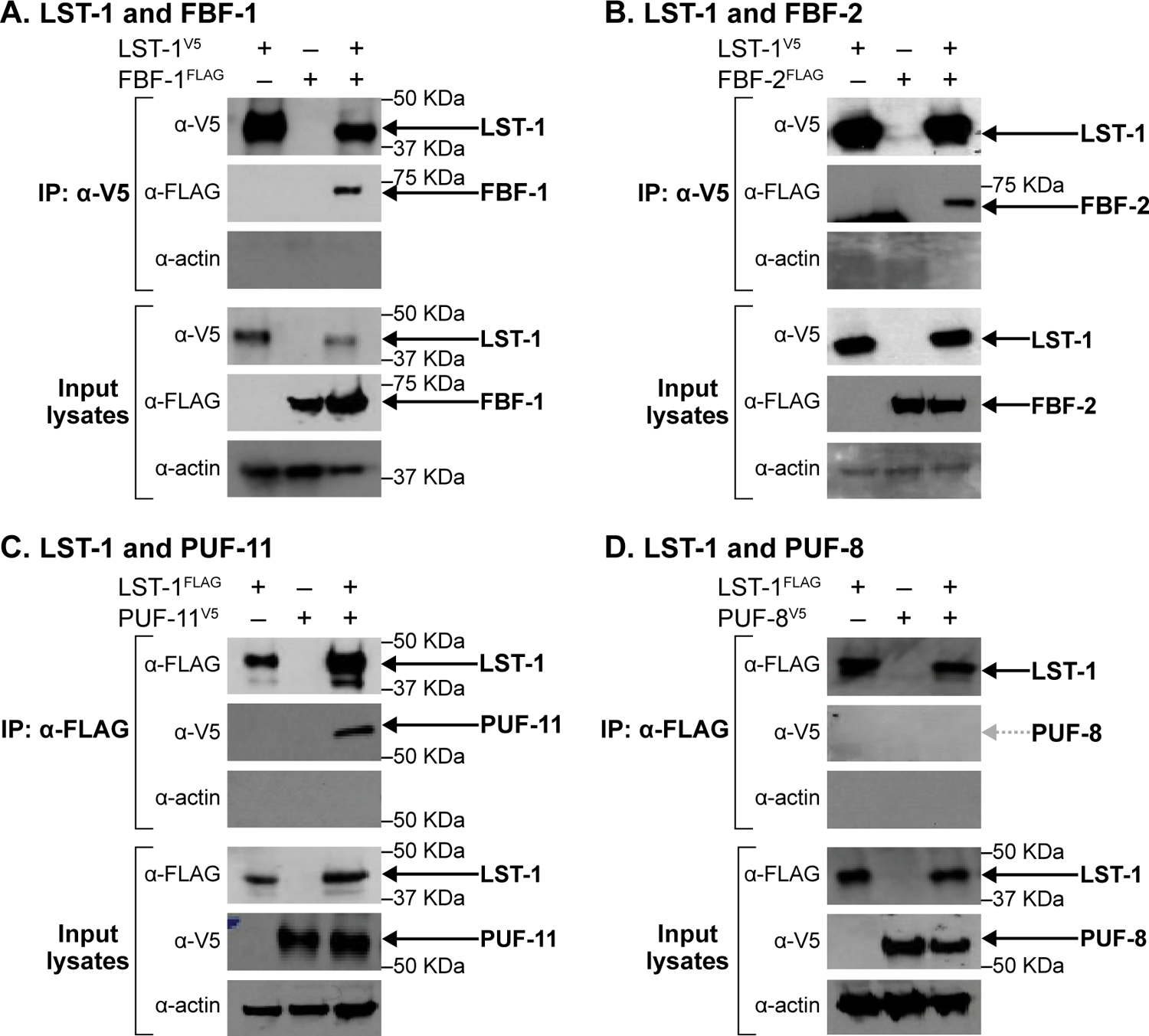
LST-1 associates specifically with PUF hub proteins in nematodes. A-D. Western blots of Input lysate and eluted samples after immunoprecipitation of epitope-tagged LST-1 from whole worms, after formaldehyde crosslinking. Blots were probed with relevant antibodies to detect epitope-tagged versions of LST-1, FBF-1 (A), FBF-2 (B), PUF-11 (C) and PUF-8 (D) as well as Actin to see the loading control. 2% of input lysates and 20% of IP-eluted samples were loaded. Exposure times were different for input and IP lanes, so band intensities are not comparable. Arrows mark LST-1 and co-immunoprecipitated proteins. Each coIP was repeated twice with similar results for the different replicates.

### ‘Kxx’mutations abrogate LST-1–FBF interactions *in vivo*

Two PUF-interacting motifs (PIMs) in LST-1 mediate its PUF interactions in yeast (see Introduction). Here we test the prediction that the PIM-defective LST-1(A^m^B^m^) mutant with 3XV5-epitope-tagged (henceforth called LST-1(A^m^B^m^) ^V5^ in the result section) would not partner with PUF proteins in nematodes. To this end, we compared the ability of LST-1 or LST-1(A^m^B^m^)^V5^ protein to immunoprecipitate FBF-1^FLAG^ or FBF-2^FLAG^ from worm lysates (Fig 3A). The protocol was as described above, with animals cross-linked prior to collection. FBF-1 and FBF-2 were abundant in input lysates, and wild-type LST-1 successfully brought down both FBF-1 and FBF-2 (Fig. 3A red box, third lane of each experiment). However, no detectable FBF-1 came down with LST-1(A^m^B^m^)^V5^ (Fig. 3A red box, fourth lane of FBF-1 experiment), and FBF-2 was sharply reduced (Fig. 3A red box, fourth lane of FBF-2 experiment). We conclude that the PIMs are indeed critical for LST-1 association with PUF proteins in nematodes and that the LST-1(A^m^B^m^)^V5^ mutant abrogates that interaction.

**Figure 3:**
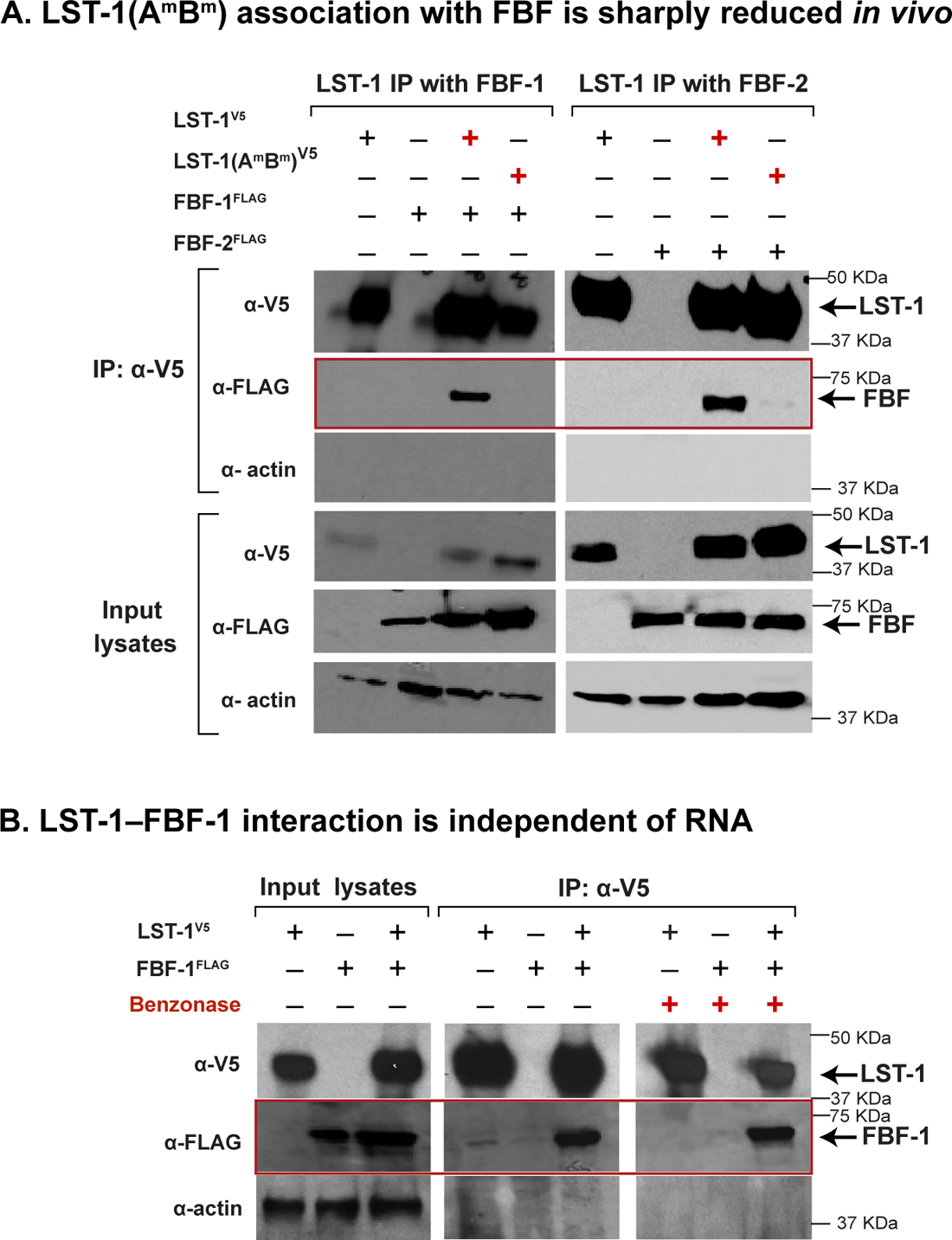
LST-1-FBF interaction is PIM-dependent and RNA-independent. A. LST-1-FBF interaction requires LST-1 PUF interaction motifs, PIM-A and PIM-B. Shown are Western blots of input lysate and eluted sample after immunoprecipitation of epitope tagged LST-1 from whole worms, after formaldehyde crosslinking. Blots were probed with α-V5 to see LST-1^V5^, α-FLAG for FBF-1^FLAG^ and FBF-2^FLAG^, and α-actin-4 for the loading control Actin. 2% of input lysates and 20% of IP-eluted samples were loaded. Each coIP was repeated at least twice with similar results for different replicates. The red box highlights presence or absence of FBFs in the LST-1 immunoprecipitate. B. LST-1-FBF interaction is independent of RNA. Shown are Western blots of input lysate and eluted sample after immunoprecipitation of LST-1^V5^ from whole worms, without formaldehyde crosslinking and with or without Benzonase. Blots were probed as described in (A). 2% of input lysates and 20% of IP-eluted samples were loaded. Each coIP was repeated twice with similar results for the different replicates. The red box highlights FBF-1 in the LST-1 immunoprecipitate.

We finally asked if the LST-1–PUF *in vivo* association depends on binding nucleic acid. In this case, worms were not subjected to formaldehyde cross-linking, and lysates were incubated prior to immunoprecipitation with Benzonase, an enzyme that cleaves both DNA and RNA. This experiment was done with worms carrying wild-type LST-1^V5^ and FBF-1^FLAG^. The efficiency of FBF-1 recovery in LST-1 IPs was similar with or without Benzonase (Figure 3B). We conclude that LST-1 associates with FBF-1 independently of both DNA and RNA and suggest that this is likely true for other LST-1–PUF partnerships.

### Tethered LST-1 represses expression of a reporter RNA

Previous experiments suggesting that LST-1 and SYGL-1 have RNA repressive activity did not examine the two proteins individually and removed them genetically, which can lead to indirect effects (Shin et al., 2017). Here, we sought to test LST-1’s regulatory activity directly and on its own. To this end, we adopted a protein-mRNA tethering strategy (Coller & Wickens, 2007). For tethering, we used the bacteriophage peptide, λN22, and BoxB sites in RNA (Baron-Benhamou et al., 2004). We introduced λN22 at the N-terminus of LST-1^V5^ and confirmed that the doubly tagged LST-1^V5-λN22^ protein is functional (Fig S3A, B). Both LST-1 tags were inserted into the endogenous gene, and LST-1^V5-λN22^ was limited as normal to the distal gonad. The reporter RNA relied on the strong germline *mex-5* promoter to drive transcription of a GFP–histone H2B RNA with three BoxB sites in its 3’UTR (Aoki et al., 2021; Aoki et al., 2018; Baron-Benhamou et al., 2004) (Figure 4A). This reporter RNA lacks a PUF binding site.

**Figure 4:**
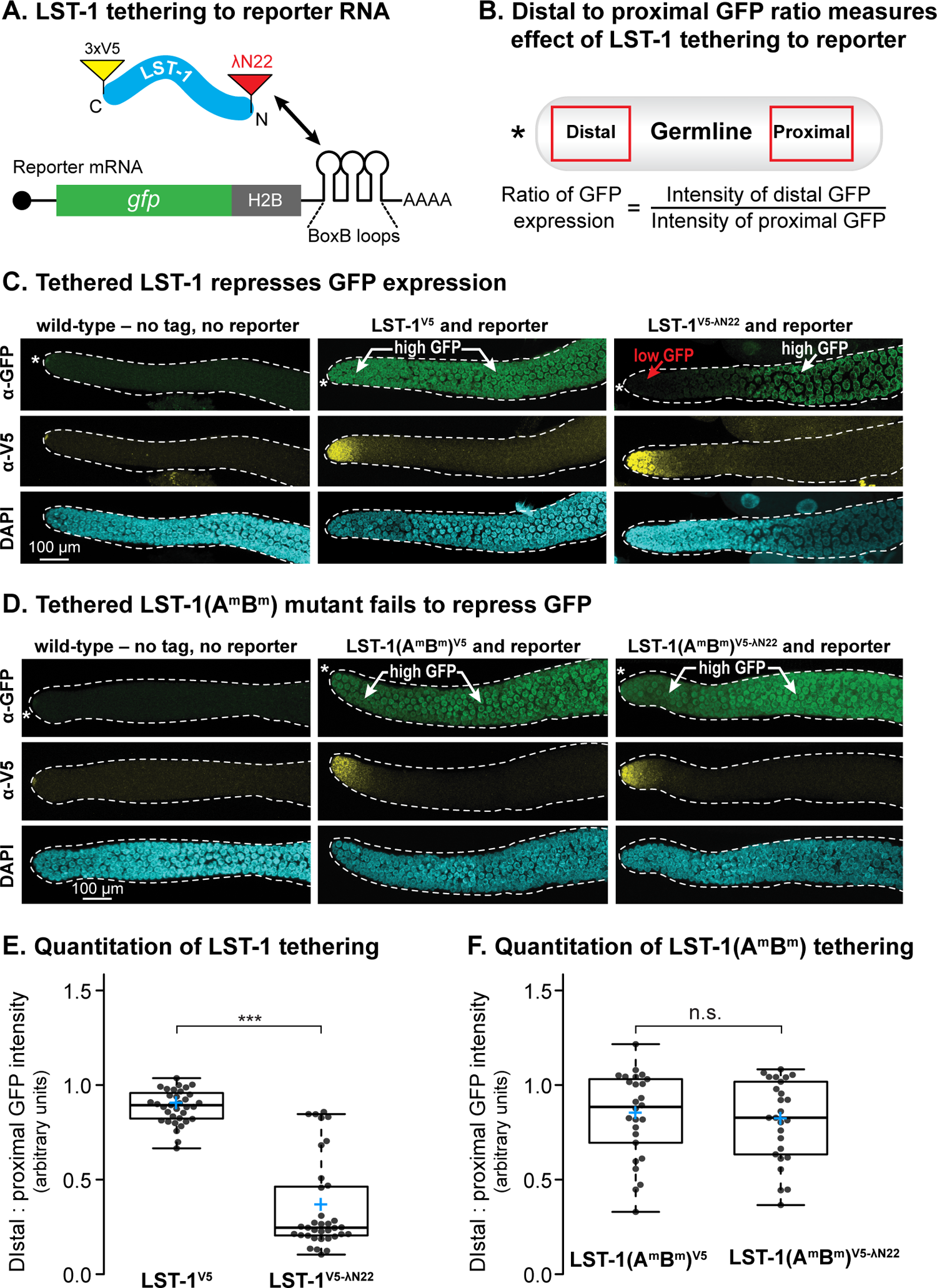
LST-1 repressive activity is PIM-dependent in tethering assay. A. Schematic of tethering assay. LST-1^V5-λN^ carries a C-terminal V5 (yellow) and N-terminal λN22 (red). LST-l^V5-λN^ binds to “Box B” hairpins for recruitment to reporter mRNA. B. Quantitating effect of tethered LST-1 on reporter expression. GFP intensity was compared in the distal germline (1-40μ from the DTC), where LST-1 is expressed at a high level, to GFP intensity more proximally (80-120μ from the DTC), where LST-1 is expressed at a vanishingly low level. C-D. Tethering results. Representative confocal images (max Z projection) of extruded gonads stained with α -GFP (top), α-V5 (middle) and DAPI (bottom). GFP is green and LST-1 is yellow when tagged; DAPI marks all gonadal nuclei. An asterisk marks the distal end. C. Tethering LST-1^V5-λN^. Left column: Control, no tag and no reporter. Middle column: Untethered LST-1, V5 and reporter but no λN22. Right column: Tethered LST-1, V5 and λN22 plus reporter. An asterisk marks the distal end. D. Tethering LST-1(A^m^B^m^)^V5-λN^ Columns same as C. LST-1(A^m^B^m^)^V5^ and LST-1(A^m^B^m^)^V5-λN^ are both restricted to distal end; LST-1(A^m^B^m^)^V5-λN^ does not repress reporter expression. Asterisk marks distal end. E. F. Boxplots of distal:proximal GFP intensity ratios. Each dot represents a separate sample. Boxes, 2575% quantile; middle line, median; blue plus sign, mean; whiskers, minimum and maximum values. Asterisks indicate statistically significant differences (student’s t.test). n.s= not significant. *** p < 0.0001. (Difference between LST-1^V5^ and LST-1^V5-λN^, p-value is 1.25 10 X^18^; difference between LST-1(A^m^B^m^)^V5^and LST-l(A^m^B^m^)^V5-λN^, p-value is 0.63). Sample sizes: LST-1^V5^, n = 35; LST-1^V5-λN^, n = 35; LST-1(A^m^B^m^)^V5^, n = 26; LST-l(A^m^B^m^)^V5-λN^, n = 26.

To measure effects of LST-1 on expression of the reporter, we compared GFP intensity in the region of the gonad where LST-1 was expressed at a high level (distal gonad) to the region where LST-1 was expressed at a vanishingly low level, just above background (proximal gonad)(Figure 4B). The ratio of distal to proximal GFP provides a quantitative measure of LST-1 RNA regulation and is internally controlled in each gonad. By GFP staining in individual gonads, the reporter was expressed in both distal and proximal regions when LST-1 was untethered (Figure 4C, middle column), but tethered LST-1^V5-λN22^ protein lowered expression distally (Figure 4C, right column). We conclude that LST-1 recruited to a reporter RNA represses expression.

We next asked if the LST-1–PUF interaction is required for repression. To this end, we inserted λN22 at the N-terminus of the LST-1(A^m^B^m^)^V5^ mutant to generate a doubly tagged LST-1(A^m^B^m^)^V5-λN22^ protein. When tested with the reporter, GFP staining was indistinguishable for untethered LST-1(A^m^B^m^)^V5 λN22^and tethered LST-1(A^m^B^m^)^V5-λN22^ proteins (Figure 4D, compare middle and right columns). Quantitation confirmed these tethering results for both wild-type LST-1^V5-λN22^ (Figure 4E) and mutant LST-1(A^m^B^m^)^V5-λN22^ (Figure 4F). We conclude that wild-type LST-1 possesses RNA repressive activity, but the mutant LST-1(A^m^B^m^)^V5^ does not. The LST-1–PUF partnership thus appears to be essential for repression.

### LST-1 associates with NTL-1 in nematode GSCs

Many RNA regulatory complexes recruit the CCR4-NOT deadenylase complex (CNOT) to repress target mRNAs (Miller & Reese, 2012; Passmore & Coller, 2022). To ask if LST-1 associates with the complex, we focused on NTL-1, the *C. elegans* homolog of the Not1 scaffold protein (DeBella et al., 2006; Nousch et al., 2013). We first inserted three tandem FLAG tags at the C-terminus of the endogenous *ntl-1* locus (Figure S4A), and confirmed that NTL-1^FLAG^ was expressed throughout the germline (Figure S4B), as seen previously for a different tagged version at the same site (Nousch et al., 2013). This NTL-1^FLAG^ protein retains biological function, as homozygous animals were viable and fertile (Figure S4C).

We first investigated the LST-1–NTL-1 association by immunostaining. LST-1^V5^ and NTL-1^FLAG^ both reside in perinuclear puncta within GSCs (Haupt et al., 2019; Nousch et al., 2013; Shin et al., 2017). Costaining revealed strong colocalization of wild-type LST-1^V5^ with NTL-1^FLAG^(Figure 5A). Most LST-1^V5^ and NTL-1^FLAG^ puncta overlapped fully or partially, while others were adjacent or did not overlap (Figure 5B). By contrast, most puncta with LST-1(A^m^B^m^)^V5^ did not overlap either fully or partially with NTL-1^FLAG^ puncta (Figure 5A, 5B). This striking difference suggests that the LST-1–NTL-1 *in vivo* association relies on the LST-1–PUF partnership.

**Figure 5:**
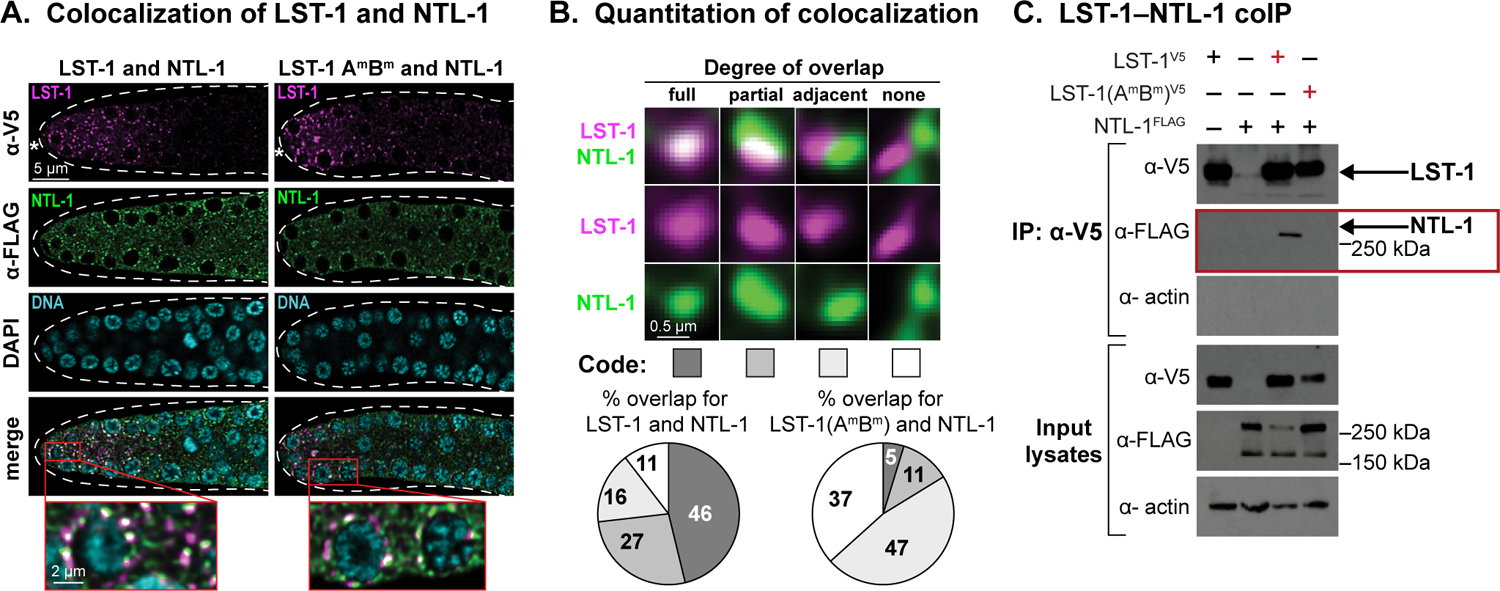
LST-1 association with NTL-1 is PIM-dependent. A. LST-1 co-localizes with NTL-1 *in vivo.* Representative deconvolved single confocal z-slices from middle plane of the distal region of an extruded gonad. Left column, strain carrying both LST-1^V5^ and NTL-1^FLAG^;right column, strain carrying both LST-1(A^m^B^m^)^V5^ and NTL-1^FLAG^. Row 1, α-V5 detects LST-1 (magenta); row 2, α -FLAG detects NTL-1 (green); row 3, DAPI highlights nuclei (cyan); row 4, merged images show co-staining with LST-1/NTL-1 overlap seen as white; insets, magnification of co-staining. Dotted line marks gonad boundary and asterisk marks distal end. B. Variable co-localization of LST-1 and NTL-1. Images show representative examples of different degrees of overlap, taken from staining in Fig 5A with further magnification. Code, shown below, is used in pie charts to show varying percentages of overlap with LST-1^V5^ (left) and LST-1 (A^m^B^m^)^V5^ mutant (right). Data in pie charts was generated from imaging 10 gonads of each strain, with 200 LST-1 foci scored in the same region of each gonad (1-30μ from distal tip). C. LST-1^V5^ and NTL-1^FLAG^ co-immunoprecipitation. Western blots were probed with α-V5 for LST-1, α-FLAG for NTL-1, and α-actin-4 for the loading control. 2% of input lysates and 20% of eluted samples were loaded. Exposure times of input and IP lanes are different, so band intensities are not comparable. The coIPs were repeated twice with similar results for the different replicates. The red box highlights presence or absence of NTL-1 in the LST-1 immunoprecipitate.

We next asked if LST-1^V5^ and NTL-1^FLAG^ co-immunoprecipitate from nematodes. The protocol was as described above, again with formaldehyde cross-linking. LST-1^V5^ did coIP with NTL-1^FLAG^ (Fig 5C red box, third lane), but LST-1(A^m^B^m^)^V5^ did not (5C red box, fourth lane). The Western blot for NTL-1 detected two major bands in the input lysates, one at ^~^260KDa and another one at ^~^170KDa, but only the larger band in the IP elutes. This larger band is similar in size to that detected previously using a LAP tag (Nousch et al., 2013). The smaller band may be a different isoform or a degradation product. The LST-1–PUF partnership thus appears to be essential for LST-1 association with the CNOT scaffold protein.

### SYGL-1 possesses two PUF-interacting motifs and has RNA repressive activity

LST-1 and SYGL-1 are functionally redundant for stem cell regulation (see Introduction), but little was known about the similarity of their molecular functions. We first asked whether SYGL-1 possesses PUF-interacting motifs. Two candidate motifs in the SYGL-1 amino acid sequence (Figure 6A) were conserved in orthologs (Figure 6B). Previous studies with other PUF partners highlighted the fourth leucine as most important (Menichelli et al., 2013; Wu et al., 2013), and crystal structures of FBF-2 with each of the LST-1 PIMs also highlighted that terminal leucine in the signature motif (Qiu et al., 2019; Qiu et al., 2022). We therefore mutated those leucines to alanines in the candidate PIMs of SYGL-1, and also mutated their N-terminal neighboring amino acid (Figure 6C). By yeast two-hybrid, wild-type SYGL-1 interacted well with both FBF-1 and FBF-2, SYGL-1 mutants defective in a single PIM lowered that interaction significantly, and mutants defective in both PIMs abolished it (Figure 6C; Figure S5C). The FBF binding strengths of the two PIMs in yeast were distinct, with PIM-A weaker than PIM-B (Figure S5C). A similar disparity was seen for the two PIMs in LST-1 (Haupt et al., 2019). Moreover, locations of the two PIMs were similar in LST-1 and SYGL-1. LST-1(PIM-A) and SGYL-1(PIM-A) begin at 32 and 39 amino acids from the N-terminus, respectively while LST-1(PIM-B) and SGYL-1(PIM-B) begin at 80 and 77 amino acids from the N-terminus, respectively. This similarity in PIM number and spacing likely relates to the geometry of their PUF binding in a way we do not yet understand. Regardless, we conclude that SYGL-1 possesses two PUF-interacting motifs.

**Figure 6:**
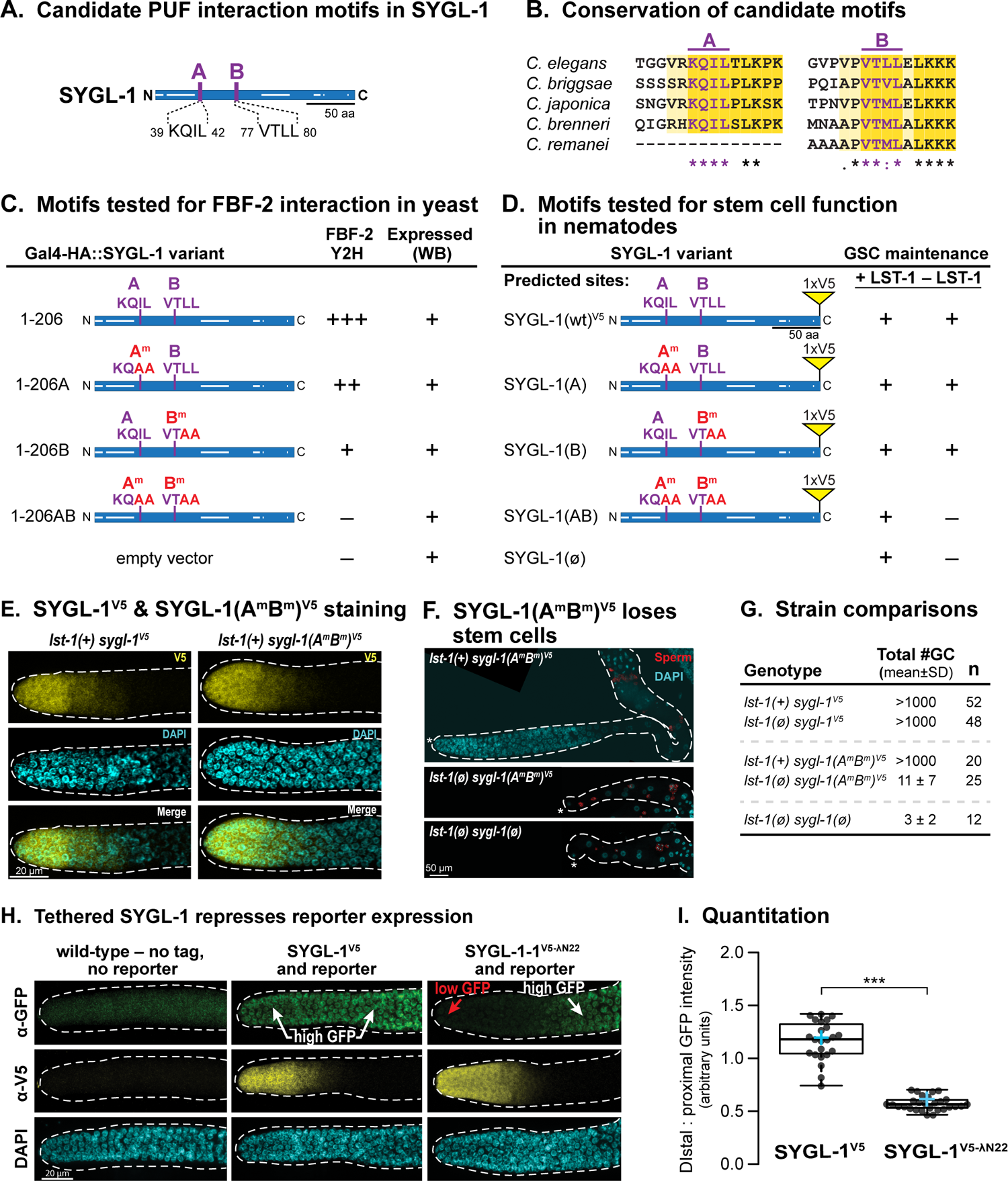
SYGL-1 has two PUF-interacting motifs and RNA repressive activity. A. Diagram of SYGL-1 with its multiple intrinsically disordered regions (white lines internal to and along the axis of the rectangle representing the protein). Two candidate PUF interaction motifs, PIM-A and PIM-B, were identified in SYGL-1 amino acid sequence. B. Conservation of PIM-A and PIM-B in SYGL-1 orthologs from related *Caenorhabditid* species. C. Summary of SYGL-1 PIM effects on FBF binding, yeast two-hybrid assay. PIM^m^ denotes a mutant, with amino acid changes in red. +++, strong binding; ++, weaker binding; +, poor binding; -, no binding. D. Summary of SYGL-1 PIM effects on GSC maintenance in nematodes. Mutation conventions as in legend to C. SYGL-1 self-renewal activity was scored both in the presence of its LST-1 redundant counterpart as a control and in the abscence of LST-1. E. Spatial restriction of SYGL-1^V5^ and SYGL-1(A^m^B^m^)^V5^ to distal gonad. Representative confocal z-projections of extruded gonads stained with α-V5 (yellow) and DAPI (cyan). An asterisk marks the distal end. F. SYGL-1(A^m^B^m^)^V5^ has lost self-renewal activity. Representative z-projected confocal images of extruded gonads stained with α-SP56 (red) for sperm and DAPI (cyan). Dotted line marks gonad boundary and asterisk marks the distal end. Top, in the presence of wild-type LST-1, SYGL-l(A^m^B^m^)^V5^ has no effect on GSC self-renewal. Middle, in absence of LST-1, SYGL-1(A^m^B^m^)^V5^ cannot maintain GSCs: the germline is tiny and GSCs differentiated in early larvae to produce a few sperm. Bottom, *Ist-1(ø) sygl-l(ø)* germlines are similar to *lst-1(ø) sygl-1(A^m^B^m^)^V5^* germlines. An asterisk marks the distal end. G. Number of total germ cells per animal in different strains. Total number of germ cells in *Ist-1(ø) sygl-1(A^m^B^m^)^V5^* is more than *lst-1(ø) sygl-1(ø)*, but fewer than *lst-1(+) sygl-1(A^m^B^m^)^V5^*. H. Tethered SYGL-1 reveals RNA repressive activity. Assay is same as Fig 4A, except λN22 is inserted at C terminus of SYGL-1^V5^. Images show representative z-projection of distal region in extruded gonads. Dotted line marks gonad boundary and asterisk marks distal end. Top, α-GFP detects GFP; middle, α-V5 detects SYGL-1; bottom, DAPI highlights DNA within gonadal nuclei. Left column: Control, no tag and no reporter. Middle column: Untethered SYGL-1, V5 and reporter but no λN22. Right column: Tethered SYGL-1, V5 and λN22 plus reporter. An asterisk marks the distal end. I. Boxplots of distal: proximal GFP intensity ratios. Conventions as described in Figure 4E, F. Asterisks indicate statistically significant differences (student t.test). *** p < 0.0001 The p value is 1.296^-14^ between SYGL-1^V5^ and SYGL-1^V5-λN^ Sample sizes: SYGL-1^V5^, n= 23; SYGL-l^V5-λN^, n= 28.

To determine whether the SYGL-1 PIMs affect stem cell function, we edited the key residues in *C. elegans* (Figure 6D). We did so in a previously edited endogenous locus that encodes a fully functional V5-tagged SYGL-1 protein. We thus generated SYGL-1(A^m^)^V5^ and SYGL-1(B^m^)^V5^ single mutants and a SYGL-1(A^m^B^m^)^V5^ double PIM mutant. All SYGL-1 variants were fertile in the presence of wild-type LST-1. In the absence of LST-1, the single PIM mutants retained their ability to maintain germline stem cells, demonstrating that one PIM is sufficient for SYGL-1 biological activity. However, the double PIM mutant was unable to maintain stem cells in the absence of LST-1 (Figure 6D), despite SYGL-1 (A^m^B^m^)^V5^ protein being expressed in GSCs (Figure 6E). In *lst-1(ø) sygl-1(A^m^B^m^)* mutants, all germline stem cells differentiated at an early larval stage (Figure 6F, 6G). We conclude that the two SYGL-1 PUF-interacting motifs are critical for stem cell maintenance.

Finally, we tested SYGL-1 for its ability to repress expression of the GFP::H2B reporter when tethered, using the assay explained above for LST-1. To this end, we inserted the λN22 peptide at the SYGL-1 C-terminus in the endogenous locus encoding wild-type SYGL-1^V5^. We thus generated a doubly tagged SYGL-l^V5-λN22^, which retains its wild-type ability to maintain stem cells in the absence of LST-1 (Figure S5D). Since SYGL-1 protein is restricted to the distal germline (Figure 6H, middle row), we quantitated its ability to repress the reporter by measuring the ratio of distal GFP to proximal GFP, as explained for LST-1 (Figure 4B). The tethered protein SYGL-1^V5-λN22^ substantially lowered GFP expression in the distal gonad (Figure 6H, 6I). Thus, SYGL-1^V5-λN22^ has repressive activity. We were unable to generate a PIM-defective SYGL-1^V5-λN22^, despite considerable effort. We conclude that SYGL-1 shares two key molecular properties with LST-1: possession of two PUF-interacting motifs essential for stem cell maintenance and ability to repress expression of an RNA when tethered.

## Discussion

### Understanding PUF hub partnerships through the lens of LST-1

The “PUF hub” of the *C. elegans* germline stem cell regulatory network provides a powerful entrée for analyzing the functional significance of partnerships between PUF RNA-binding proteins and their modulating partners (see Introduction, Figure 1A). Here, we test key elements of the PUF hub model in nematodes for the first time and investigate how the PUF partnerships regulate germline stem cells. We focus on LST-1–PUF as a paradigm, because a nematode mutant had been created with potential to assess the *in vivo* function of LST-1–PUF partnerships. This LST-1(A^m^B^m^) mutant lacks amino acid residues responsible for its PUF interaction in yeast (Kimberly A Haupt et al. 2019b). We confirm in this work that wild-type LST-1 associates with PUF hub proteins in nematodes, but that the LST-1(A^m^B^m^) mutant protein does not. Therefore, assembly of LST-1–PUF partnerships in nematodes depends on PUF-interacting motifs (PIMS), as predicted. However, the LST-1–PUF complex did not depend on RNA. Because each PIM can act independently to bind PUFs in yeast and to promote GSC self-renewal in worms (Kimberly A Haupt et al. 2019b; Qiu et al. 2019; Qiu et al. 2022), our model for LST-1–PUF assembly includes two distinct complexes, one anchored by PIM-A (Fig. 7a, left) and the other by PIM-B (Fig. 7a, right), but not an RNA. We conclude that LST-1 forms PIM-dependent but RNA-independent partnerships with PUF hub proteins in the nematode (Figure 7A).

**Figure 7:**
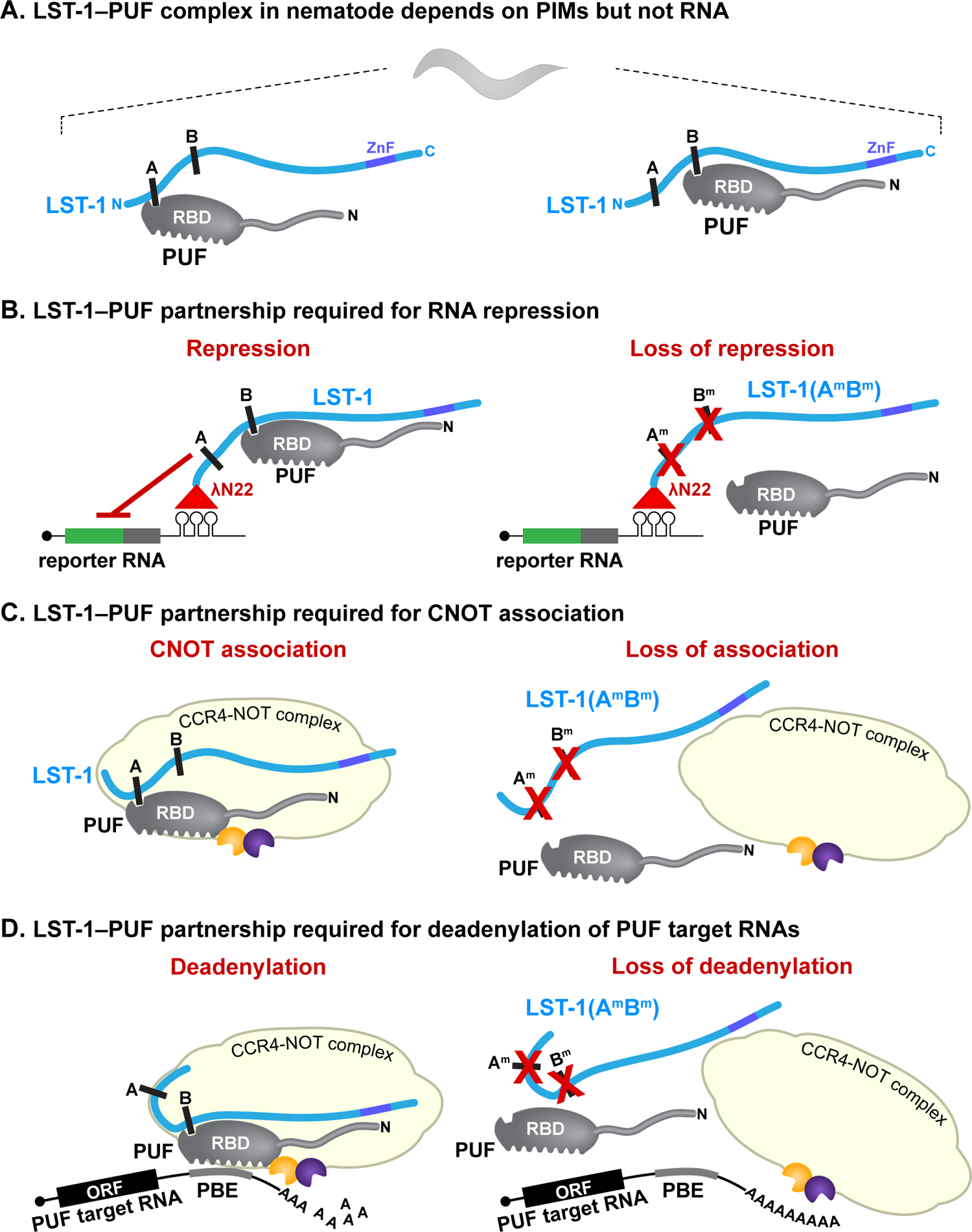
Models for assembly and function of LST-1–PUF partnership in nematodes. A. Model: Assembly of LST-1–PUF partnerships. LST-1 (blue) can bind to PUF proteins (grey) via either of two PUF-interacting motifs (A and B) (Haupt et al., 2019). No RNA is depicted because LST-1–PUF assembly does not require RNA (this work). PUF proteins comprise an RNA binding domain (RBD) and an N-terminal tail (wavy line) with intrinsically disordered regions (IDRs)(Fig S6). LST-1 is largely intrinsically disordered (wavy line) and also has C-terminal Zinc finger (purple); LST-1 stem cell function resides in its IDR region and does not require the Zn finger. Fig 7A conventions are same in Fig 7B-D models. B. Model: LST-1 RNA repressive activity depends on PUF partnership. Left, when tethered, the LST-1–PUF complex represses expression of the reporter RNA; a PUF protein is included in this diagram, because PIM-defective LST-1(A^m^B^m^) cannot repress reporter RNA expression when tethered (right). Tethering employs λN22 (red triangle) fused to LST-1 to bind boxB stem loops in reporter RNA (see Fig 4A). C. Model: LST-1-CNOT association depends on PUF partnership. Left, LST-1–PUF complex associates with CNOT complex (light yellow); right, PIM-defective LST-1(A^m^B^m^) mutant disrupts LST-1–PUF complex and destabilizes CNOT association. Dark yellow and blue pacman figures, CCF-1 and CCR-4 deadenylases. The model includes an interaction between the PUF protein and CCF-1, based on work with FBF-1 and FBF-2 (Suh et al., 2009) D. Model: LST-1–PUF partnership brings together multiple interactions to form a stable complex with the CNOT complex and repress target RNAs. Left, LST-1–PUF complex provides LST-1 IDRs and PUF IDRs; PUF protein contacts CCF-1 and also binds PUF binding element (PBE) in its target RNA. Right, PIM-defective LST-1(A^m^B^m^) mutant disrupts LST-1–PUF and destabilizes the larger complex (LST-1–PUF–CNOT–target RNA), shown here as loss of all interactions for simplicity.

Previous experiments implicated LST-1 and SYGL-1 in RNA repression (Shin et al. 2017). However, those experiments did not test the two proteins separately; they did not test significance of the PUF partnership; and they did not identify the likely effector of RNA repression. This work tackles all three issues and does so in nematode GSCs – the natural context. We used a tethering assay to investigate LST-1 RNA regulatory activity. This assay is direct, and it queries LST-1 separately from SYGL-1. The tethered LST-1 dramatically lowered expression of the reporter RNA. A common interpretation of this result would be that LST-1 acts alone to repress RNA. However, we also tethered the LST-1(A^m^B^m^) mutant, which cannot assemble an LST-1–PUF complex. To our surprise, LST-1(A^m^B^m^) lost RNA repressive activity. Although LST-1 PIMs might mediate binding to some non-PUF protein, the PIM-dependent PUF interaction is remarkably specific (self-renewal PUF proteins only), and we favor the simpler explanation – that LST-1 retains its partnerships with PUF proteins when recruited to the reporter RNA (Figure 7B). How might an LST-PUF partnership repress RNA? PUF proteins recruit the CNOT complex to their target RNAs in virtually all eukaryotes – from yeast and plants to flies and humans (Nishanth & Simon, 2020). Nematode FBF-1 and FBF-2, for example, interact physically with a subunit of the CNOT complex, the nematode homolog of the CAF1 deadenylase, and they also promote its enzymatic activity *in vitro* (Suh et al., 2009). Here we show that LST-1 co-localizes in subcellular puncta with the nematode homolog of the Not1 scaffold protein of the CNOT complex, called NTL-1 (Nousch et al., 2013), and in addition, that LST-1 coIPs with NTL-1. However, the PIM-defective LST-1(A^m^B^m^) mutant dramatically reduces co-localization and does not coIP with NTL-1, suggesting that LST-1 repressive activity and LST-1–NTL-1 association both depend on the LST-1–PUF partnership (Fig 7C).

Why is the LST-1–PUF partnership critical for RNA repression and association with CNOT? Answering that question in molecular detail will require future experiments to analyze formation of a larger LST-1–PUF–CNOT–RNA complex and map the key interaction surfaces. However, this work together with results from others suggests the model that LST-1–PUF provides multiple interactions that work together to form a stable effector complex on PUF target RNAs (Fig 7D). One type of interaction relies on regions predicted to be intrinsically disordered (IDRs). In yeast, fly and human PUF proteins, IDRs associate with CNOT and enhance its deadenylase activity (Arvola et al., 2020; Enwerem et al., 2021; Webster et al., 2019) with longer IDRs enhancing better than shorter ones (Webster et al., 2019). The PUF IDRs are located in N-terminal “tails” – variably long extensions N-terminal to the PUF RNA-binding domain. The fly Pumilio N-terminal tail harbors an IDR-rich RD3 domain that interacts specifically with the Not1, Not2 and Not3 subunits of the CNOT complex (Haugen et al., 2022). Nematode PUF proteins also have IDR-rich N-terminal tails (Fig. S6), but the tails are short (^~^120 amino acids in worms versus ^~^1000 in flies and ^~^800 in humans). The LST-1–PUF partnership therefore brings a PUF protein with its N-terminal IDRs and LST-1 with its IDRs in the “self-renewal region” (Fig 1B) into a single complex (Fig. 7A). It seems likely that PUF and LST-1 IDRs work together to facilitate association with CNOT (Fig 7D, left). A second interaction, seen *in vitro*, occurs between the FBF-1 and FBF-2 RNA binding region and one CNOT subunit, the CCF-1/CAF deadenylase enzyme (Suh et al., 2009). FBF-1 and FBF-2 physically interact with CCF-1 and promote its deadenylase activity. A third interaction occurs between the PUF protein and its PUF binding element in the target RNA. We suggest that these interactions together form a stable effector complex that promotes deadenylation and represses target RNAs (Fig 7D, left). We also suggest that the effector complex is destabilized without the LST-1–PUF complex (Fig 7D, right), with the diagram suggesting loss of all interactions for simplicity.

### SYGL-1 and LST-1 share molecular features critical for PUF hub function

SYGL-1 and LST-1 are redundant for stem cell maintenance, but their amino acid sequences bear no similarity to each other. However, both consist largely of intrinsically disordered regions. Although LST-1 has a C-terminal Zinc Finger, its stem cell function resides in the IDRs, not the Zinc finger (Haupt et al., 2019). A key question has been whether the IDRs of SYGL-1 and LST-1 employ similar molecular mechanisms. This work identifies two common molecular features, suggesting that they do.

First, SYGL-1 and LST-1 both possess two PUF-interacting motifs (PIMs) that function independently to maintain stem cells. Thus, SYGL-1 retains its self-renewal capacity when either PIM is mutated, but loses it when both are defective; the same is true of the two LST-1 PIMs (Haupt et al., 2019, this work). We do not know why LST-1 and SYGL-1 have two PIMs when a single PIM is sufficient for stem cell maintenance. In yeast two-hybrid assays and *in vitro*, the two LST-1 PIMs differ in their FBF binding affinity, with PIM-A weaker than PIM-B (Haupt et al., 2019; Qiu et al., 2019; Qiu et al., 2022); the same is true of the two SYGL-1 PIMs in yeast (this work). However, those different affinities may not be relevant in nematodes if mitigated by multiple interactions of the LST-1–PUF partnership with the CNOT complex. Previous work discovered a KTxL signature for PIMs in CPB-1 and GLD-3 (Menichelli et al., 2013; Wu et al., 2013). Identification of the LST-1 PIMs revealed a related KxxL sequence (Haupt et al., 2019; Qiu et al., 2019), which also was found in SYGL-1 PIM-A (this work). The SYGL-1 PIM-B sequence (VTLL), however, reduces the consensus PIM motif to a single leucine (this work). That leucine is critical for PUF binding in a spectrum of *in vitro* and *in vivo* assays, but additional molecular features must exist to provide context for its binding (Qiu et al., 2022). With the range of PIM sequences now available and others likely to emerge, we can ask if all FBF partners have two PIMs and *how in vitro* PIM differences affect *in vivo* function.

The second molecular feature common to LST-1 and SYGL-1 is their RNA repressive activity when tethered. PUF proteins are well known for RNA repression, and above we discuss how the LST-1–PUF complex is critical for repression of target mRNAs (Fig 7D). Here, we suggest that the SYGL-1–PUF complex likely uses the same mechanism to recruit CNOT to PUF target mRNAs and repress them.

### LST-1–PUF and Nos-Pum partnerships: similarities and differences

*Drosophila* Nanos and Pumilio provide a well-established paradigm for interaction of a PUF protein with its partner (Arvola et al., 2017). Our growing understanding of the *C. elegans* LST-1–PUF partnerships invites comparison. The fly Nos-Pum and worm LST-1–PUF complexes share several features. Both regulate germline stem cells (Forbes & Lehmann, 1998; Shin et al., 2017); both repress RNAs (Sonoda & Wharton, 1999; Wang et al., 2020; this work); and both recruit the CNOT complex (Kadyrova et al., 2007; this work). In addition, Nanos and LST-1 both interact with their respective PUF proteins in the same region – at the loop between 7^th^ and 8^th^ repeats of the RNA-binding domain (Qiu et al., 2019; Qiu et al., 2022; Weidmann et al., 2016). And finally, spatial restriction of both Nanos and LST-1 is responsible for localizing PUF function to a specific region. Nanos localization in the posterior embryo restricts PUF-dependent RNA repression to that region, and LST-1 localization to the distal gonad restricts PUF-dependent RNA repression to germline stem cell pool. Therefore, one might think *a priori that* LST-1 and SYGL-1 would be analogous to Nanos and that their PUF partnerships would function similarly.

Yet that simple idea seems to be wrong. One difference is obvious from their distinct use of Zinc fingers. The two Zinc Fingers in Nanos are integral to its ternary complex with Pumilio and RNA (Curtis et al., 1997; Lehmann & Nusslein-Volhard, 1991; Weidmann et al., 2016). These Nanos Zinc Fingers bind RNA just upstream of the Pumilio response element and contribute to a molecular clamp that strengthens Pumilio binding to RNA (Weidmann et al., 2016). By contrast, the single Nanos-like Zinc Finger in LST-1 can be deleted without affecting LST-1–PUF regulation of stem cells (Haupt et al., 2019). Furthermore, SYGL-1, the redundant counterpart of LST-1, does not possess a Zinc finger, underscoring the irrelevance of the Zinc finger to the partnership. A second difference emerges from biochemical experiments testing how the partners affects PUF affinity for RNA. Nanos enhances PUF affinity for RNA (Weidmann et al., 2016), but PIM-bearing fragments of LST-1 weaken it (Qiu et al., 2019; Qiu et al., 2022). This LST-1 conclusion may be misleading, however, given its reliance on a peptide with a single PIM. It will be important to learn whether full length LST-1 strengthens or weakens PUF affinity for RNA. A third difference is that Nanos cannot bind stably to Pumilio without RNA (Arvola et al., 2017; Sonoda & Wharton, 1999; Weidmann et al., 2016), but LST-1 binds to FBF independently of RNA (this work). And finally, tethered Nanos represses RNA on its own (Raisch et al., 2016), but LST-1 requires its PUF partnership (this work). Thus, the Nos-Pum and LST-1–PUF complexes represent two distinct modes of PUF partnership, despite their both having a repressive RNA regulatory activity that relies on the CNOT complex. Those distinct modes showcase the LST-1–PUF partnership as an emerging paradigm for understanding how PUF proteins are modulated by partners to control gene expression.

## Acknowledgements

The authors thank members of the Kimble and Wickens labs for helpful discussions throughout the course of this work. We thank Jane Selegue, Peggy Kroll-Conner, and Sadie Jackson for assistance with experiments or strain building. We thank Traci M Tanaka Hall and Chen Qiu for critical comments on the manuscript and Laura Vanderploeg for help with figures. This work was supported by Kamaluddin Ahmad Distinguished Graduate Scholarship to ASF; National Science foundation Graduate Research Fellowship Program under Grant Nos. DGE-1256259 and DGE-1747503 to BHC; NIH R01 GM50942 to MW; and NIH R01 GM134119 to JK. (Any opinions, findings, and conclusions or recommendations expressed in this material and those of the authors(S) and do not necessarily reflect the views of the National Science Foundation).

## Methods and Materials

### Strain Maintenance

*C. elegans* strains were maintained as described (Brenner, 1974) at 20°C except those with *glp-1(gf ts)*, which were maintained at 15°C but shifted to 20°C or 25°C for experimentation. For a complete list of strains used in this study, see Table S1. Strains are available upon request.

### RNA interference

RNA interference (RNAi) was performed by feeding, as described (Kershner et al., 2014; Timmons & Fire, 1998). *sygl-1* and *lst-1* clones from the Ahringer RNAi library (Fraser et al., 2000) were used for RNAi treatment, and L4440 plasmid (‘empty’ RNAi) for the negative control. Detailed protocol can be found in (Kershner et al., 2014).

### CRISPR Cas9 induced allele generation

CRISPR–Cas9 genome editing methods were used to alter endogenous *lst-1, sygl-1* and *ntl-1* genes (Table S1) using ribonucleoprotein complexes with a co-conversion strategy following an established protocol (Arribere et al., 2014; Paix et al., 2015). For detailed protocol, see (Haupt et al., 2019). Sequences of all crRNA (IDT) and DNA templates (IDT) are listed in table S2.

### Scoring tumor production using DAPI staining

Germline tumors induced by *glp-1(ts gf)* at 25° were scored with DAPI (4□,6-diamidino-2-phenylindole) staining of extruded gonads, following the protocol described by (Crittenden et al., 2017), with some modifications. Briefly, animals were dissected in PBStw [PBS+0.1% (v/v) Tween-20] with 0.25 mM levamisole to extrude gonads, then fixed at room temperature for at least 15 min in 2% paraformaldehyde diluted in PBStw. Samples were incubated overnight at −20°C in 100% methanol. Next day the samples were washed with PBStw, then incubated with 0.5 ng/μl DAPI in PBStw to label DNA. Then, samples were mounted in either Vectashield (Vector Laboratories) or ProLong Gold (Thermo Fisher Scientific). Tumors were confirmed by observation of proliferation throughout the germline, including metaphase plates proximally, and few if any gametes. Some germlines had patches of meiotic cells as previously described (Pepper et al., 2003).

### Immunostaining, Microscopy, Fluorescence quantitation

Immunostaining was performed as described (Crittenden et al., 2017) with minor modifications. Briefly, animals were staged to 24 h past mid-L4 stage (when grown at 20°C) or 18 hr past mid-L4 stage (when grown at 25°C). Staged animals were dissected in PBStw (PBS + 0.1% (v/v) Tween-20) with 0.25 mM levamisole to extrude gonads after cutting behind pharynx. Tissues were fixed in 3% (w/v) paraformaldehyde diluted in 100 mM K_2_HPO_4_ (pH 7.2) for 20 min. After fixation, all samples were permeabilized with ice-cold methanol (for worms that harbor GFP) for 20 min or PBStw + 0.2% (v/v) Triton-X for 5–10 min. Samples were then washed twice by adding PBSTw followed by centrifugation at 1500 rpm for 60s, then excess liquid was removed. Next, they were blocked with either 30% (v/v) goat serum diluted in PBStw (for anti-FLAG) or 0.5% (w/v) bovine serum albumin diluted in PBStw (all other antibodies) for 1 h. Primary antibodies were then added and samples incubated overnight at 4°C in blocking solution at the following dilutions: mouse *α*-FLAG (1:1000, M2 clone, Sigma #F3165), mouse *α*-V5 (SV5-Pk1, 1:1000, MCA1360, Bio-Rad) and mouse *α*-SP56 (1:200, a gift from Susan Strome, University of California, Santa Cruz, CA, USA). Samples were washed twice the next day with PBSTw. For secondary antibodies, samples were incubated for 1 h at room temperature in dark at the following dilutions: Donkey Alexa 555 *α*-mouse (1:1000, Invitrogen #A31570), Donkey Alexa 647 *α*-mouse (1:500, Invitrogen #A31571). To visualize DNA, DAPI was added at 0.5–1 ng/μl during the last 20 min of secondary antibody incubation. Samples were then washed twice with PBSTw to remove excess antibodies. After the last wash, excess liquid was removed, and samples were mounted in ProLong Gold (Thermo Fisher Scientific) on microscope slides (Cat #12-544-1, Fisher Scientific) and covered with 22×22 coverslip (Fisher scientific different supplier). Mounted samples were cured overnight before imaging. All images were taken using a laser scanning Leica TCS SP8 confocal microscope with LASX software. Photomultiplier (PMT) detectors were used for DAPI and Hybrid (HyD) detectors were used for all other fluors. A 63X/1.40 CS2 HC Plan Apochromat oil immersion objective was used for all images, which were taken with the standard 400-700 Hz scanning speed and 100-300% zoom. Immunostaining quantitation was performed using Fiji/ImageJ. For detailed protocol, see (Haupt et al., 2019)

For the tethering assays, GFP intensity in the distal germline (1-40μm from distal end) was compared to that more proximally (80-120μm from the DTC micron) in the same germline using FIJI/ImageJ. Ratios of distal to proximal intensity were calculated using Microsoft excel software. Samples from at least three independent replicates were analyzed together after normalizing to a control with no GFP (N2).

### Progenitor zone count

Progenitor zone (PZ) size was scored in DAPI-stained extruded gonads from hermaphrodites 24 h past mid-L4 at 20°C or 18 h past mid-L4 at 25°C. PZ sizes were scored following the convention described in (Crittenden et al., 2006), (Seidel & Kimble, 2015). Scoring was done manually using the FIJI/ImageJ multi-point tool; each DAPI stained nucleus along the edge of the tissue was considered a unique cell row; values from the two edges of the gonad were then averaged to determine PZ size.

### Immunoprecipitations and Western blotting

*glp-1(gf ts)* animals were raised at 15°C or 20°C until they became gravid; they were then bleached to obtain synchronized offspring. Synchronized L1 worms were put at 25°C for 48 hours to induce germline tumors. A minimum of 10^6^ young adults were collected using the following protocol: animals were washed twice with M9 buffer [3 g/L KH2PO4, 6 g/L NaHPO4, 5 g/L NaCL, and 1 mM MgSO4] and cross-linked with 1% (w/v) formaldehyde for 10 min at room temperature. For immunoprecipitations in Figure 3B, samples were not cross-linked before collection. Worm pellets were then washed twice with M9 and snap frozen in liquid nitrogen for subsequent analysis. Pellets were resuspended in 1 ml lysis buffer [20 mM Tris pH 7.5, 150 mM NaCl, 2 mM EDTA, 5mM MgCl_2_, 1% (v/v) Triton-X, 1 M Urea, complete Protease inhibitor cocktail (Roche)]. Urea was not used for samples processed to produce Figure 3B. Worms were lysed by adding 1 sterilized Retsch 5-mm stainless steel ball to each sample (is this correct?), and then put in a Retsch 400 MM mill mixer at 4°C for three 10-minute cycles at 30 Hz. After cycle 1 and 2, two 5-minute freeze-thaw steps were done by immersion in liquid nitrogen for 1 min followed by immersion in room temperature water for 4 min. Lysates were cleared twice by centrifugation (16,000g, 15 min at 4°C), and the total protein concentration was measured using Bradford assay (Biorad).

To prepare antibody conjugated beads, 20 μg mouse *α*-V5 (Bio-Rad #MCA1360) (For Figure 2A, 2B, 3A, 3B) or mouse *α*-FLAG (M2 clone, Sigma #F3165) (For Figure 2C and 2D) was incubated with 4.5 mg protein G Dynabeads (Novex, Life Technologies, #10003D) for 60 minutes at RT. The Dynabeads were then washed to remove unbound antibodies. The total amount of protein for IP was normalized to input. 20 mg lysates were incubated with the antibody-bead mixture for 4 h at 4°C, in the presence of RNase A at 10 μg/ml. RNA degradation was confirmed by isolating total RNA from post-IP lysates using TRIzol LS (Invitrogen #10296028) and analyzing on agarose gels. For IP in Figure 3B, 1 μL of Benzonase^®^ Nuclease (Sigma milipore, 250U/uL) was used instead of RNase A for specific samples (see Figure 3, legend). Beads were pelleted, washed four times with lysis buffer, and then two times with wash buffer [20 mM Tris pH 7.5, 0.5 M NaCl, 2 mM EDTA, 1% (v/v) Triton X-100]. Samples then were eluted with elution buffer [1% (w/v) SDS, 250 mM NaCl, 1 mM EDTA, 10 mM TRIS pH 8] for 10 min at 100°C and analyzed by Western blotting.

For Western blotting, input and eluted samples were run on 10% acrylamide gel at room temperature. Then samples were transferred to PVDF membrane (Immobilon-P, 0.45 μm Merck Millipore Ltd.), which was activated prior to transferring in 100% methanol following a wash in ddH2O. The transfer was carried out for 4 h at 4°C in transfer buffer containing 20% Methanol. For NTL-1 (Figure 5), the transfer buffer contained 10% methanol to minimize precipitation of the large protein. After transfer, the membrane was blocked for 1 h at RT in 5% skimmed powdered milk. For primary antibodies, blots were incubated overnight at 4°C at the following dilutions: Mouse *α*-FLAG (1:1000, M2 clone, Sigma #F3165), Mouse *α*-V5 (1:1000, Bio-Rad #MCA1360), Mouse *α*-actin (1:40,000, C4 clone, Millipore #MAB1501). For secondary antibodies, blots were incubated for 1 h at RT with Rat HRP-conjugated *α*-mouse (1:10,000, Abcam mAb 131368). To analyze the co-immunoprecipitations, blots were stripped with Restore^™^ Western Blot Stripping Buffer (Thermo scientific). Immunoblots were developed using SuperSignal^™^ West Pico/Femto Sensitivity substrate (Thermo Scientific #34080, #34095) and imaged using an ImageQuant LAS4000 (GE Healthcare). FIJI/Image J was used to adjust contrast. For each set of samples Co-IPs were done at least twice.

### Co-localization assay

Co-localization (Figure 5) was scored manually using FIJI/ImageJ. Confocal images of extruded gonads stained for DNA and both epitope-tagged LST-1 and NTL-1 were processed using the Leica Lightning deconvolution package. Composite images were used to score co-localization of LST-1 and NTL-1. LST-1 foci were first marked using multi-point tool in the distal 30 μm of the gonad using only the LST-1 channel. The NTL-1 channel was then added, and the degree of colocalization was manually scored. 200 LST-1 foci were scored in each gonad. The experiment was repeated twice with a total of 20 gonads for each strain.

### Yeast two-hybrid

Modified yeast two-hybrid assays were performed as described (Bartel & Fields, 1997). Briefly, *sygl-1* cDNA encoding wild-type full-length SYGL-1 (a.a. 1-206), or full-length SYGL-1 carrying PIM mutations were cloned into the Nco I site in pACTII (Gal4 activation domain plasmid), generating pJK1580, pJK1581, pJK1582, pJK2094, pJK2095, and pJK2096 respectively, using the Gibson assembly method. The PUF repeat region of FBF-2 (a.a. 121-632) was cloned into the Nde I site in pBTMknDB (LexA binding domain plasmid) to generate pJK2046. Activation and binding domain plasmids were co-transformed into the L40-ura strain using the Te-LiAc method (Gietz & Schiestl, 2007). LacZ reporter activity was assayed in defined media (SD) supplemented with -Leu-Trp using the Beta-Glow^®^ Assay System following commercially available protocols (Promega #E4720). In short, yeast cultures were grown to mid-log phase, diluted to the same optical density (0.1), and added to equal volumes of Beta-Glow^®^ reagent. Yeast clones were then incubated for 1 hr at room temperature, and luminescence was quantitated using a Biotek Synergy 4 Hybrid plate reader and Gen5 software (Winooski, VT). A complete list of plasmids used in yeast two-hybrid assays is available in Table S3.

### Statistical Analysis

Statistical analyses, sample sizes and *p* values are described in figure legends. A two-tailed Student’s ř.test (T.TEST function in Microsoft excel) assuming equal-variance was done when comparing two samples. A p-value less than 0.05 was considered significant. Box plots were generated with web tool BoxPlotR (Spitzer et al., 2014), quartiles and whiskers are indicated in legends.

