## Supplemental Fig 1 for "Functional significance of PUF partnerships in *C. elegans* germline stem cells"

**A LST-1$^{FLAG}$ distribution in *glp-1(gf ts)* gonads**

**20ºC - permissive** **25ºC -- restrictive**

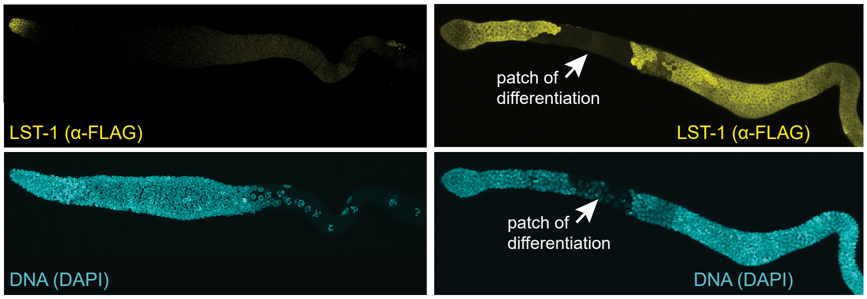


**B LST-1(A$^m$B$^m$)$^{V5}$ distribution in *glp-1(gf ts)* gonads that also carry FBF-2$^{FLAG}$**

**20ºC - permissive** **25ºC -- restrictive**

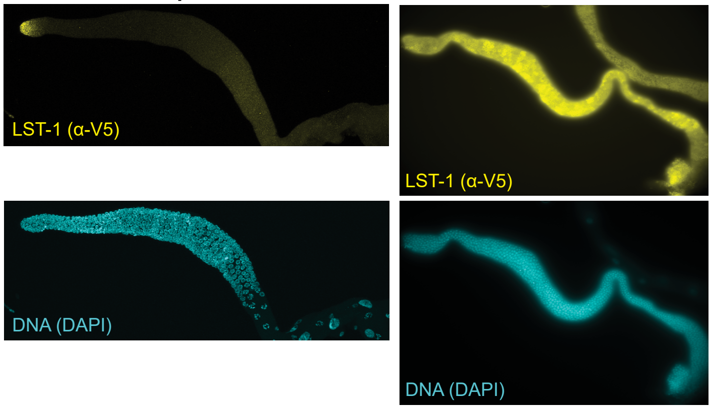


**C. Phenotype characterization**

| Genotype* | % Fertility 20°C | n | PZ size mean gcd ± sd 20°C | n | % Tumor 25°C | n |
|---|---|---|---|---|---|---|
| *lst-1$^{V5}$* | 100 | 100 | 19±1 | 10 | 0 | 50 |
| *lst-1$^{V5}$ ; glp-1(gf ts)* | 98 | 100 | 20±3 | 10 | 100 | 50 |
| *lst-1(A$^m$B$^m$)$^{V5}$* | 97 | 100 | 19±2 | 10 | 0 | 50 |
| *lst-1(A$^m$B$^m$)$^{V5}$ ; fbf-1$^{FLAG}$; glp-1(gf ts)* | 98 | 100 | 22±1 | 10 | 100 | 50 |
| *lst-1(A$^m$B$^m$)$^{V5}$ ; fbf-2$^{FLAG}$; glp-1(gf ts)* | 95 | 100 | 19±3 | 10 | 99 | 50 |

* See Table S1 for allele and strain numbers.
For Progenitor zone (PZ) size, n is number of gonadal arms scored.
