## Supplemental Fig 2 for "Functional significance of PUF partnerships in *C. elegans* germline stem cells"

Figure S2
Ferdous et al 2022

### A Phenotype characterization

| Genotype * | % Fertility 20°C | n† | PZ size mean gcd ± sd 20°C | n† | % Tumor 25°C | n† | Reference |
|---|---|---|---|---|---|---|---|
| *fbf-1$^{FLAG}$* | 100 | 112 | 19±2 | 10 | 0 | 100 | this work |
| *fbf-1$^{FLAG}$ fbf-2(ø)* | 70§ | 133 | 24 ± 3 | 11 | nd | n/a | this work |
| *fbf-1$^{FLAG}$; glp-1(gf ts)* | 98 | 100 | 21±1 | 15 | 100 | 50 | this work |
| *lst-1$^{V5}$; fbf-1$^{FLAG}$ ; glp-1(gf ts)* | 100 | 100 | 20±2 | 15 | 100 | 50 | this work |
| *fbf-2$^{FLAG}$* | 100 | 62 | 20±1 | 62 | 0 | 50 | this work |
| *fbf-1(ø) fbf-2$^{FLAG}$* | 100 | 53 | 15±2 | 53 | n/a | n/a | this work |
| *fbf-2$^{FLAG}$; glp-1(gf ts)* | 100 | 100 | 19±3 | 15 | 100 | 50 | this work |
| *lst-1$^{V5}$; fbf-2$^{FLAG}$; glp-1(gf ts)* | 100 | 100 | 21±1 | 15 | 100 | 50 | this work |
| *puf-11$^{V5}$* | 100 | 100 | 18±2 | 15 | 0 | 100 | this work |
| *puf-11$^{V5}$; glp-1 (gf ts)* | 100 | 100 | 21±2 | 15 | 100 | 50 | this work |
| *lst-1$^{FLAG}$; puf-11$^{V5}$; glp-1(gf ts)* | 100 | 100 | 19±1 | 15 | 100 | 50 | this work |
| *puf-8$^{V5}$* | 100 | 100 | 20±2 | 15 | 0 | 100 | this work |
| *puf-8$^{V5}$; glp-1(gf ts)* | 100 | 100 | 22±1 | 25 | 100 | 50 | this work |
| *llst-1$^{FLAG}$; puf-8$^{V5}$; glp-1(gf ts)* | 100 | 100 | 20±1 | 25 | 100 | 50 | this work |

* See Table S1 for allele and strain numbers.

† For % Fertile and % Tumor, n is number animals scored; for Progenitor zone (PZ) size, n is number of gonadal arms scored.

§ Non-fertile animals of this genotype make normal-sized germlines that are Fog (no sperm, oocytes only).

### B Distribution of PUF proteins in *glp-1(gf ts)* worms at restrictive and permissive temperature

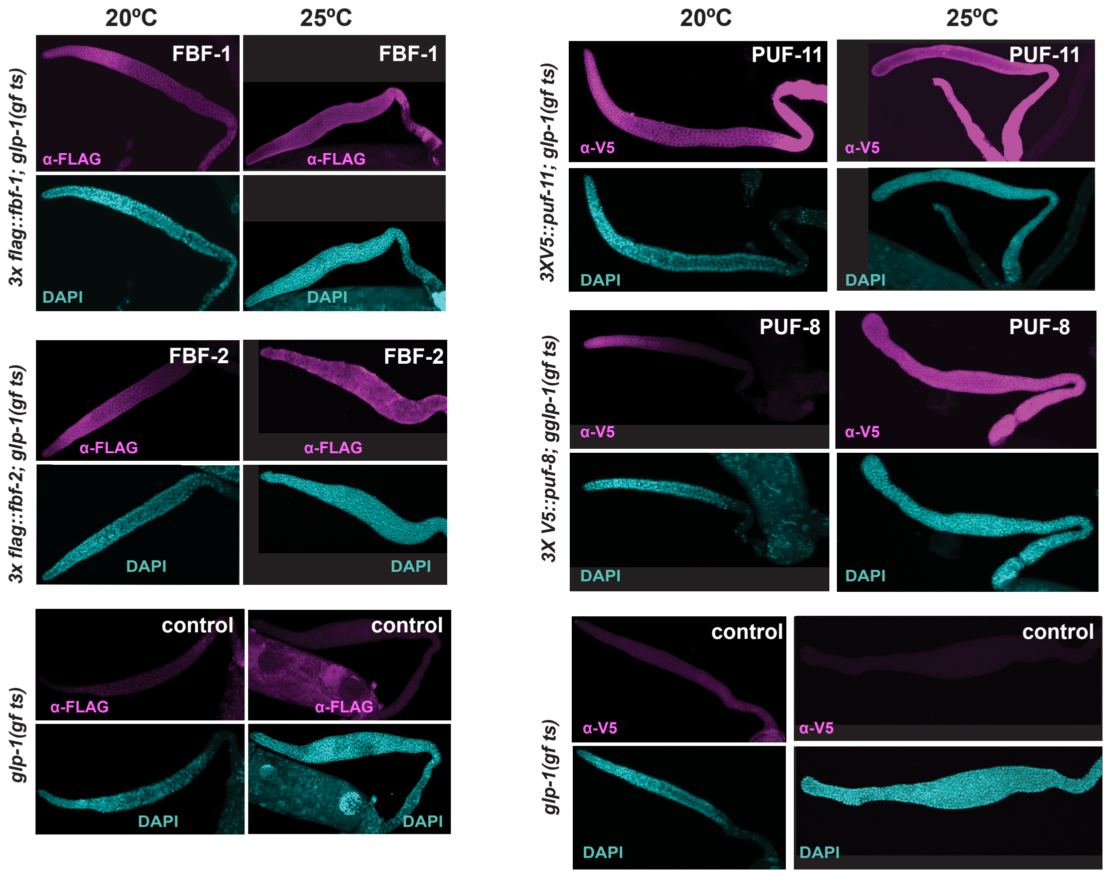
