## Supplemental Fig 3 for "Functional significance of PUF partnerships in *C. elegans* germline stem cells"

**A Comparison of LST-1$^{V5}$ and LST-1$^{V5\text{-}\lambda N22}$ expression**

**B Comparison of LST-1($A^mB^m$)$^{V5}$ and LST-1($A^mB^m$)$^{V5\text{-}\lambda N22}$ expression**

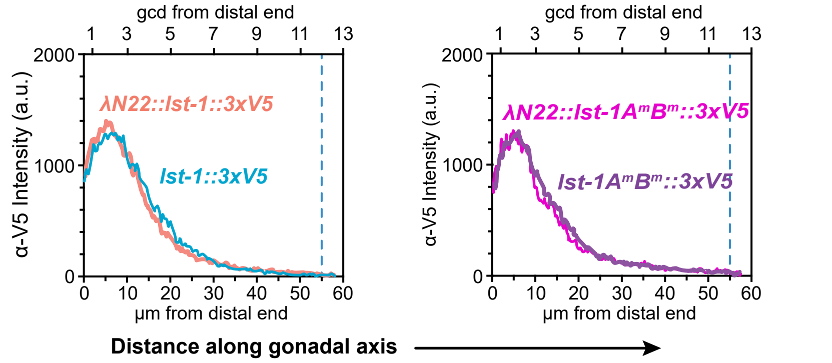


### C LST-1$^{V5\text{-}\lambda N22}$ retains biological function

| Genotype* lst-1 | sygl-1 | % Fertile 20°C | n† | PZ size mean gcd ± sd 20°C | n† |
|---|---|---|---|---|---|
| *lst-1$^{V5}$* | + | 99 | >100 | 20.0±2.0 | 10 |
| *lst-1$^{V5}$* | ø | 97 | >100 | 13.2±1.5 | 15 |
| *lst-1$^{V5\text{-}\lambda N22}$* | + | 96 | 100 | 19.6±2.5 | 15 |
| *lst-1$^{V5\text{-}\lambda N22}$* | ø | 97 | 100 | 13.0±2.0 | 15 |
| *lst-1($A^mB^m$)$^{V5}$* | + | 98 | >100 | 20.0±1.0 | 15 |
| *lst-1($A^mB^m$)$^{V5}$* | ø | 0 | 100 | n/a | 0 |
| *lst-1($A^mB^m$)$^{V5\text{-}\lambda N22}$* | + | 98 | 100 | 21.0±2.0 | 15 |
| *lst-1(ø)* | RNAi | 0 | 100 | n/a | 0 |

* See Table S1 for allele and strain numbers.

† For % Fertile, n is number animals scored;
for Progenitor zone (PZ) size, n is number of gonadal arms scored.
