## Supplemental Fig 4 for "Functional significance of PUF partnerships in *C. elegans* germline stem cells"

**A. Epitope tagging the endogenous *ntl-1* gene locus**

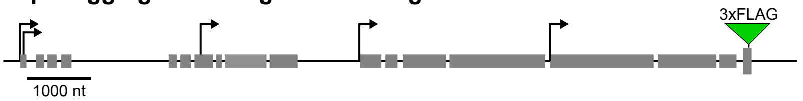


**B. NTL-1 protein distribution in germline tumor**

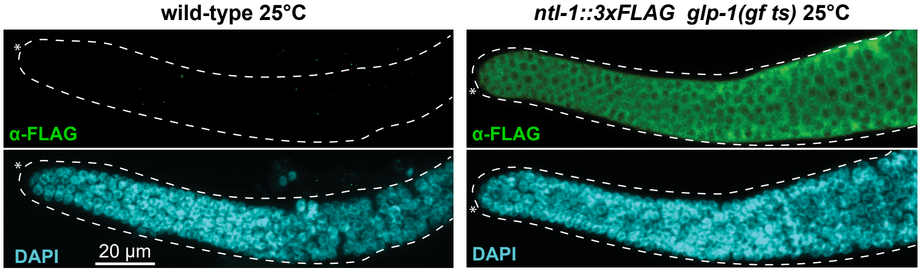


**C. Phenotype characterization**

| Genotype* | % Fertility 20°C | n | PZ size mean gcd ± sd 20°C | n | Brood size mean ± sd 20°C | n^ | %Tumor 25°C | n |
|---|---|---|---|---|---|---|---|---|
| wild-type | 98 | 100 | 20±1 | 21 | 211±15 | 4 | 0 | 100 |
| *ntl-1*$^{FLAG}$ | 99 | 100 | 19±2 | 25 | 220±13 | 4 | 0 | 100 |
| *ntl-1*$^{FLAG}$ *glp-1(gf ts)* | 99 | 100 | 19±1 | 25 | 215±15 | 4 | 100 | 100 |
| *lst-1*$^{V5}$; *ntl-1*$^{FLAG}$ *glp-1(gf ts)* | 97 | 100 | 19±2 | 30 | 220±10 | 4 | 100 | 100 |
| *lst-1(A*$^{m}$*B*$^{m}$*)*$^{V5}$; *ntl-1*$^{FLAG}$ *glp-1 (gf ts)* | 97 | 100 | 20±2 | 20 | 210±10 | 4 | 100 | 100 |

* See Table S1 for allele and strain numbers. For Progenitor zone (PZ) size, n is number of gonadal arms scored.
^number of F0 worms, F1 generations from these worms were scored for brood size;
