## Supplemental Fig 5 for "Functional significance of PUF partnerships in *C. elegans* germline stem cells"

**A. Two-hybrid assay** **B. β-galactosidase assay** **C. Western blot**

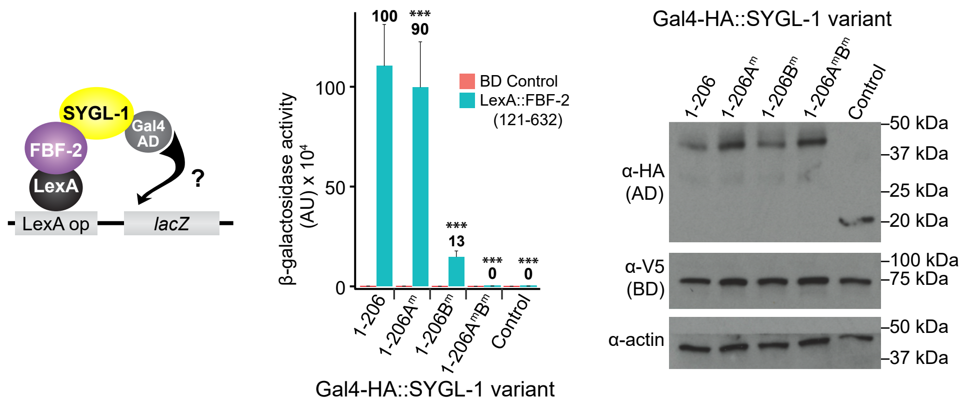


### D. Both SYGL$^{V5}$ and SYGL-1$^{V5-\lambda N22}$ function normally to maintain germline stem cells

| Genotype | % Fertility | n | # PZ Size mean gcd ± sd | n |
|---|---|---|---|---|
| *lst-1(+) sygl-1(+)* | 98 | 100 | 20 ± 2 | 20 |
| *lst-1(ø) sygl-1$^{V5}$* | 95 | 60 | 20 ± 1 | 20 |
| *lst-1(ø) sygl-1$^{V5-\lambda N22}$* | 94 | 81 | 19 ± 3 | 20 |
| *lst-1(ø) sygl-1(ø)* | 0 | 100 | n/a | 10 |

For Progenitor zone (PZ) size, n is number of gonadal arms scored.
