## Supplementary figures and images for "Functional significance of PUF partnerships in *C. elegans* germline stem cells"

### Supplemental Fig 6

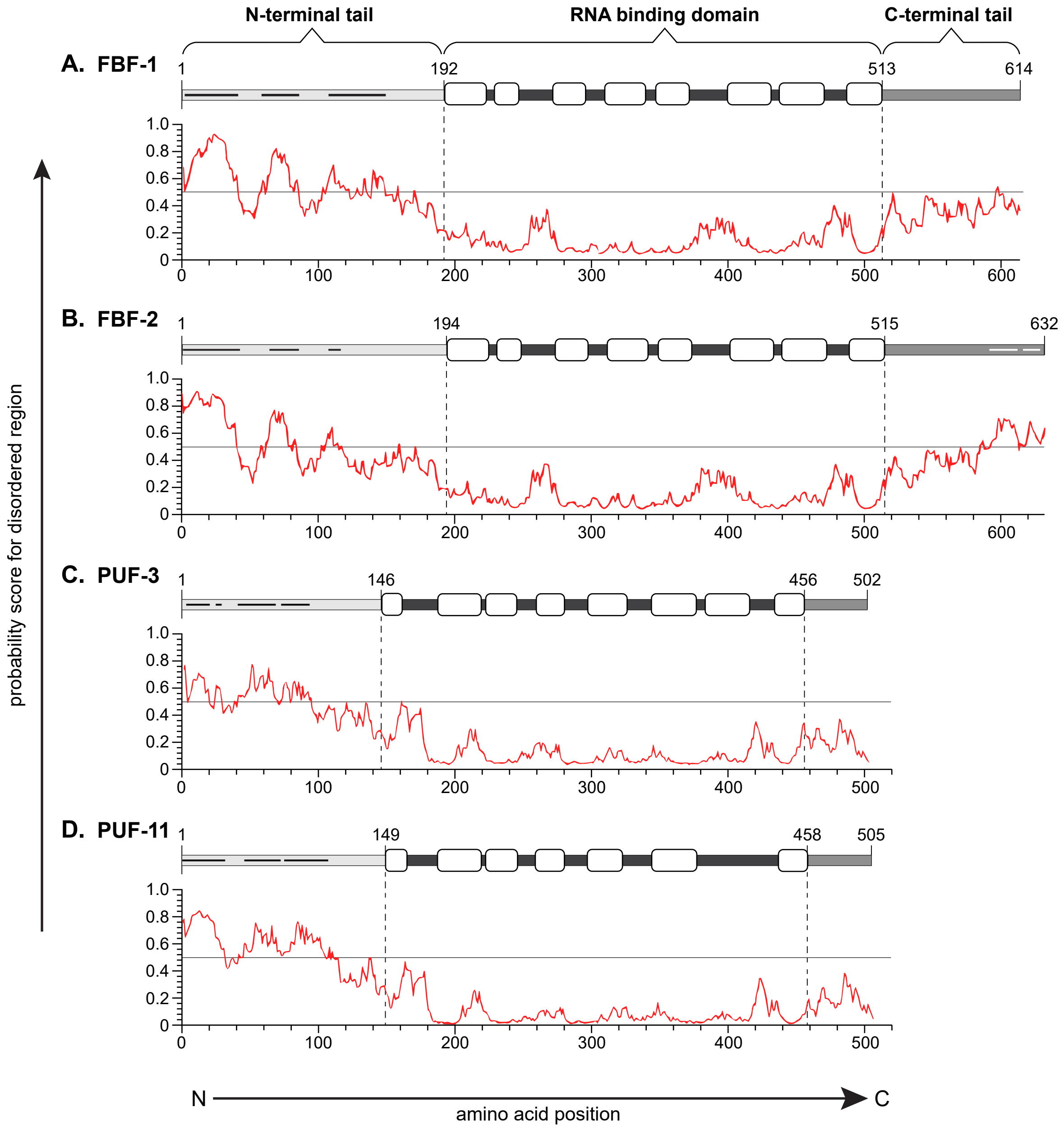
