## Supplemental Fig legends and tables for "Functional significance of PUF partnerships in *C. elegans* germline stem cells"

Supplemental figure legends

**Figure S1.** Supplementary analysis of LST-1 and LST-1(A^m^B^m^ ) proteins.
A. Representative z-projection of extruded gonads showing LST-1^FLAG^ distribution at permissive and restrictive temperature in *glp-1 (gf ts)* animals.

B. Representative z-projection of extruded gonads showing LST-1(A^m^B^m^ )^V5^ distribution at permissive and restrictive temperature in *glp-1 (gf ts)* background.

C. Phenotypic characterization of lst-1(A^m^B^m^ ) mutants. Fertility refers to self-fertility. PZ size refers to progenitor zone size, measured as the mean number of germ cell diameters (gcd) from the distal end to meiotic entry. The mean PZ size for wild-type is of 20 germ cell diameter (Crittenden et al., 2006). Tumor refers to a germline tumor.

**Figure S2.** Supplementary analysis of tagged PUF proteins.
A. Phenotypic characterization of animals carrying PUF proteins that were epitope tagged at the endogenous locus. Conventions as in Fig S1C.

B. Representative z-projection of extruded gonads showing distribution of PUF proteins
at permissive temperature (20°C) and restrictive temperature (25°C).

**Figure S3:** Supplementary analysis of LST-1^V5-λN22^ protein.
A. Expression: Left, LST-1^V5^ and LST-1^V5-λN22^ are expressed with similar abundance along axis of extruded gonads stained with α-V5. Right, LST-1(A^m^B^m^ )^V5^ and LST-1(A^m^B^m^ )^V5-λN22^ are expressed with similar abundance along axis of extruded gonads stained with α-V5.

B. Self-renewal activity: Conventions as in Fig S1C; brood size refers to total number of self-progeny produced by a single hermaphrodite. LST-1^V5^ and LST-1^V5-λN22^ both retain self-renewal activity; LST-1(A^m^B^m^)^V5^ and LST-1(A^m^B^m^)^V5-λN22^ both lose self-renewal activity.

**Figure S4 :** Supplementary analysis of NTL-1^FLAG^.
A. Diagram of *ntl-1* gene shown to scale with all its alternative transcription start sites. The FLAG tag at the NTL-1 C-terminus labels all NTL isoforms.
B. Expression: Left, untagged NTL-1 is not detected; right, NTL-1^FLAG^ is expressed throughout the germline. Representative z-projected confocal images of extruded gonads stained using 𝛼-FLAG to detect NTL-1^FLAG^ (green) and DAPI to detect nuclei (cyan).
C. Phenotype: Conventions as in Fig S1C. At 20 °C, NTL-1^FLAG^ has no effect on Progenitor Zone Size, Brood Size or Fertility; at 25°C, NTL-1^FLAG^ has no effect on tumor formation.

**Figure S5.** Supplementary SYGL-1 experiments

A. Schematic of yeast two-hybrid assay to test SYGL-1–FBF-2 interaction. Variants of full length SYGL-1(1-206) protein were fused to Gal4 activation domain (AD), and FBF-2(121-632), which includes all PUF repeats, was fused to LexA binding domain (BD). Interaction drives transcription of lacZ genes.

B. Results of β-gal assay. Each bar is an average of at least three independent replicates; error
bars show standard deviation. Activity for each variant is indicated as a percentage of the activity of wild-type SYGL-1, with the number above each bar. Asterisks indicate a statistically significant difference from wild-type by one-way ANOVA with Tukey’s HSD test. ***, p<0.001; n.s., not significant (p >0.05).

C. Expression Gal4 fusion variants, assayed by Western blot.

D. Phenotype: Conventions as in Fig S1C. SYGL-1^V5-λN^ retains self-renewal activity.

**Figure S6:** Predicted intrinsically disordered regions in *C. elegans* PUF proteins specialized for self-renewal.

A-D: Top of each panel, schematic of PUF protein according to PFAM database, with its N-terminal tail (light grey), RNA binding domain with eight PUF repeats (rounded rectangles), and C-terminal tail (dark grey). Lines internal to and along the axis of the protein schematic mark intrinsically disordered regions. Bottom of each panel, probability of intrinsic disorder (red line) according to IUpred3 (using long disorder algorithm with default setting, (Erdős et al., 2021). A probability score of ≥0.5 (horizontal grey line) indicates disordered regions in all N-terminal tails and the C-terminal tail of FBF-2; a probability score of ≤0.5 indicates order regions in all RNA binding domains. Red lines were rendered from the IUpred3 original file in illustrator by image trace

**Supplementary tables**

**Table S1: Nematode strains^1^ used in this study.**

| Strain | *Genotype* | *Refer to in the text* | | *Reference* |
| --- | --- | --- | --- | --- |
| N2 | *Wild type* | *wild-type* | *(*Brenner, 1974*)* | |
| JK5629 | *glp-1(ar202) [G529E] III/ hT2[qIs48](I;III)* | *Stock* | *(*Pepper et al., 2003*)* | |
|  | *glp-1(ar202)* | *glp-1(gf ts)* |  | |
| JK5758 | *lst-1(q895) [lst-1::3XFLAG] I* | *LST-1^FLAG^* | *(*Haupt et al., 2019*)* | |
| JK5810 | *fbf-2(q945) [3xFLAG fbf-2] II* | *FBF-2^FLAG^* | *This work* | |
| JK5929 | *lst-1(q1004) [lst-1::3xV5] I* | *LST-1^V5^* | *(*Shin et al., 2017*)* | |
| JK5935 | *fbf-1(ok91) [fbf-1 deletion] fbf-2(q945) [3xFLAG fbf-2] II* | *FBF-1 Ø FBF-2^FLAG^* | *This work* | |
| JK6002 | *sygl-1(q1015) [sygl-1::1XV5] I* | *SYGL-1^V5^* | *(*Shin et al., 2017*)* | |
| JK6004 | *sygl-1(q1017) [sygl-1::1xV5::GGS linker::λN] I* | *SYGL-1^V5-λN22^* | *This work* | |
| JK6026 | *lst-1(q1028) [λN::lst-1::3xV5] I* | *LST-1^V5-λN22^* | *This work* | |
| *JK6063* | *puf-8(q1048) [3xV5::puf-8] II* | *PUF-8^V5^* | *This work* | |
| JK6203 | *lst-1(q1125) [(L35A, K80A, L83A)::3xV5] I* | *LST-1 (A^m^B^m^) ^V5^* | *(*Haupt et al., 2019*)* | |
| JK6270 | *puf-11(q1128) [puf-11::3xV5] IV* | *PUF-11^V5^* | *(*Haupt et al., 2020*)* | |
| JK6319 | *lst-1(q1004) [LST-1::3xV5] sygl-1(q828) [sygl-1 deletion] I* | *LST-1^V5^ sygl-1(ø)* | *(*Shin et al., 2017*)* | |
| JK6332 | *lst-1(q1004) I; glp-1(ar202) III* | *LST-1^V5^; glp-1(gf ts)* | *This work* | |
| JK6367 | *qSi375[(mex-5 promoter::eGFP::linker::his-58::3xboxb::tbb-2 3’utr) *weSi2] II* | *tethering GFP reporter* | *(*Aoki et al., 2021*)* | |
| JK6371 | *sygl-1(q1188)*: *[(I41A, L42A, L79A, L80A) sygl-1::1xV5]* | *SYGL-1(A^m^B^m^) ^V5^* | | *This work* |
| JK6372 | *lst-1(q1125) I; fbf-2(q945) II; glp-1(ar202) III* | *LST-1 (A^m^B^m^) ^V5^; FBF-2^FLAG^; glp-1(gf ts)* | | *This work* |
| JK6373 | *fbf-2(q945) II; glp-1(ar202) III* | *FBF-2^FLAG^ glp-1(gf ts)* | | *This work* |
| JK6374 | *lst-1(q1004) I; fbf-2(q945) II; glp-1(ar202) III* | *LST-1^V5^; FBF-2^FLAG^; glp-1(gf ts)* | | *This work* |
| JK6385 | *sygl-1(q1162) [(I41A, L42A) sygl-1::1xV5] I* | *SYGL-1 (A^m^) ^V5^* | | *This work* |
| JK6386 | *sygl-1(q1189) [(L79A, L80A) sygl-1::1xV5] I* | *SYGL_1 (B^m^) ^V5^* | | *This work* |
| JK6494 | *lst-1(q895) I; glp-1 (ar202) III* | *LST-1^FLAG^ glp-1(gf ts)* | | *This work* |
| JK6524 | *ntl-1 (q1236) [ntl-1::3XFLAG] III* | *NTL-1^FLAG^* | | *This work* |
| JK6537 | *fbf-1(q1241) [3x FLAG:FBF-1] II* | *FBF-1^FLAG^* | | *This work* |
| JK6571 | *fbf-1 (q1248) [3x FLAG:FBF-1] fbf-2 (q738)[deletion of puf repeats]* | *FBF-1^FLAG^ FBF-2 Ø* | | *This work* |
| JK6579 | *puf-8(q1048) II; glp-1(ar202) III* | *PUF-8^V5^ ; glp-1(gf ts)* | | *This work* |
| JK6580 | *lst-1(q1004) I; fbf-1(q1241) II; glp-1(ar202) III* | *LST-1^V5^; FBF-1^FLAG^; glp-1(gf ts)* | | *This work* |
| JK6581 | *fbf-1(q1241) II; glp-1(ar202 gf) III* | *FBF-1^FLAG^ ; glp-1(gf ts)* | | *This work* |
| JK6582 | *lst-1(q895) I; glp-1(ar202) III; puf-11(q1128) IV* | *LST-1^FLAG^; glp-1(gf ts); PUF-11^V5^* | | *This work* |
| JK6583 | *glp-1(ar202) III; puf-11(q1128) IV* | *PUF-11^V5^ glp-1(gf ts)* | | *This work* |
| JK6584 | *ntl-1(q1244) [3XFLAG] glp-1(ar202) III* | *NTL-1^FLAG^ glp-1 (gf ts)* | | *This work* |
| JK6618 | *lst-1(q1265) [lst-1(λN: L35A, K80A, L83A)::3xV5] I* | *LST-1(A^m^B^m^)^V5-λN22^* | | *This work* |
| JK6641 | *lst-1(q1004) I; qSi375II* | *LST-1^V5^; tethering reporter* | | *This work* |
| JK6642 | *lst-1(q1028) I; qSi375II* | *LST-1^V5-λN22^; tethering reporter* | | *This work* |
| JK6643 | *lst-1(q1125) I; qSi375 II* | *LST-1 ( A^m^B^m^ ) ^V5^; tethering reporter* | | *This work* |
| JK6644 | *lst-1(q1265) I; qSi375 II* | *LST-1( A^m^B^m^ )^V5-λN22^; tethering reporter* | | *This work* |
| JK6696 | *lst-1(q1004) I; ntl-1(q1244) glp-1(ar202) III* | *LST-1^V5^ ; NTL-1^FLAG^ glp-1 (gf ts)* | | *This work* |
| JK6697 | *lst-1(q1125) I; ntl-1(q1244) [3XFLAG] glp-1(ar202) III* | *LST-1( A^m^B^m^ ) ^V5^; NTL-1^FLAG^ glp-1 (gf ts)* | | *This work* |
| JK6698 | *lst-1(q1004) sygl-1(q828) I; glp-1(ar202) III* | *LST-1^V5^ sygl-1(ø); glp-1 (gf ts)* | | *This work* |
| JK6700 | *lst-1(q1125) I; fbf-1(q1241) II; glp-1(ar202 gf) III* | *LST-1 ( A^m^B^m^ ) ^V5^; FBF-1^FLAG^; glp-1(gf ts)* | | *This work* |

^1^Strains are available upon request.

**Table S2: CRISPR guide RNA and repair oligos used in this work.**

| gene | strain | guide RNA sequence (5'-3') | DNA repair oligo sequence (5'-3')^1^ |
| --- | --- | --- | --- |
| *puf-8* | *q1048* | uugaaaucggacgacucauc | ctaaagtaattattttaggggtaatctgcgttcctgATGggtaagcctatccctaaccctctcctcggtctAgatAGTacTggAaagccAatcccAaacccActcctcggActtgatAGCacCggtaagcctatccctaacccActcctcggActtgatAGCacCTCAAGACCAATTTCAATTGGGAATACATGCACATT |
| *fbf-1* | *q1241* | aagcccacatctacagaggt | atgctgcgccatcggcgaatgtgcgggagtatgatgatttccCTTGTCATCGTCATCCTTGTAATCGATGTCATGATCTTTATAATCACCGTCATGGTCTTTGTAGTCaacctctgtagatgtgggcttttgcggagtctgtaaaattta |
| *fbf-2* | *q945* | gatctttataatcaccgtca | aatattattttcatattcccaatcattctaataaaattatcaactaatcgacATGGACTACAAAGACCATGACGGTGATTATAAAGATCATGACATCGATTACAAGGATGACGATGACAAGGATCAATCTAAAATGCGAAGAACAAATCAGTTCAGAAAAgtaagttttcagtaacttctaaagaaacaaatggcctaag |
| *ntl-1* | *q1244* | accagatgagaaaaatcagt | aataccggagctgctaatcaacagcaaaacccaaacacAaacGACTACAAAGAtCATGACGGTGATTATAAAGATCATGACATCGATTACAAGGATGACGATGACAAGtgatttttctcatctggtctttcggttcccgattttctgatc |
| *lst-1* | *q1028, q1265* | aaaagtgaactcactcttgg **agg** | caaaagaattatttgactattcaatcttcgcgcgagacaATGGGAAACGCCCGTACCCGTCGTCGTGAGCGTCGTGCCGAGAAGCAAGCCCAATGGAAGGCCGCCAACGGAGGATCCTGTTCGACTTTTATTCTTCCTCCTCGTgtgagttcactttttttttgaaattaaatatgtattttctct |
| *sygl-1* | *q1162* | aatctgcttgcggacgcctc | cgccacgtcatcatcgtcaacaggaggcgtccgcGCCGCCacgttgaagccaaagcaaccgaatggttatgtgcaacact |
| *sygl-1* | *q1188* | caccggaaccgggacaccgc | atcttcggtgatttccggcggtgtcccggttccggtgacTGCCGCCgagttgaagaagaaggatcgtagcgagaagaatgtggtga |
| *sygl-1* | *q1189* | caccggaaccgggacaccgc | agtgttgcacataaccattcggttgctttggcttcaacgtGAGAATctgcttgcggacgcctcctgttgacgatgatgacgtggcg |
| *sygl-1* | *q1017* | cuacugcaaauaauagcuguguuuuagagcuaugc | gaacaacaacacttcactgatgatgggctcTaacagctattatttgcagggtaagcctatccctaaccctctcctcggtctagatagtactGGAGGATCCGGAAACGCCCGTACCCGTCGTCGTGAGCGTCGTGCCGAGAAGCAAGCCCAATGGAAGGCCGCCAACtagagcgtacttgctcttttaaattttctaatcc |

^1^Uppercase letters denote mutations (including insertions, PAM mutations and/or seed

sequence mutations)

| Plasmid | Insert Description | Cloning Site | Vector Backbone | Reference |
| --- | --- | --- | --- | --- |
| pJK1580 | HA::SYGL-1 (a.a. 1-206) | Nco I | pACT2 | (Shin et al., 2017) |
| pJK1581 | HA::SYGL-1 (a.a. 1-103) | Nco I | pACT2 | This work |
| pJK1582 | HA::SYGL-1 (a.a. 103-206) | Nco I | pACT2 | This work |
| pJK2094 | HA::SYGL-1 (a.a. 1-206), A Site | Nco I | pACT2 | This work |
| pJK2095 | HA::SYGL-1 (a.a. 1-206), B Site | Nco I | pACT2 | This work |
| pJK2096 | HA::SYGL-1 (a.a. 1-206), AB Site | Nco I | pACT2 | This work |
| pJK2046 | V5::FBF-2 (a.a. 121-632) | Nde I | pBTMKnDB | (Haupt et al., 2019) |

**Table-S3: Plasmids used in this work for yeast two-hybrid assay**
